# Tau aggregation results in an impaired ability to counteract microtubule destabilization via specific combinatorial tau phosphorylation patterns

**DOI:** 10.1101/2025.08.09.669485

**Authors:** Nima N Naseri, Tian Xu, Ibrahim Saleh, Sapanna Chantarawong, Steven M Patrie, Neil L Kelleher, Manu Sharma, Hong Xu, Edward B. Lee, Ophir Shalem, Elizabeth Rhoades

**Author notes:** **CORRESPONDING AUTHOR** Correspondence to Elizabeth Rhoades.

## Abstract

Tau phosphorylation is a defining feature of Alzheimer’s disease, yet it also plays an important physiological role in regulating functional interactions with microtubules (MTs), particularly during normal neuronal development. While individual tau phosphorylation sites have been well-studied, how combinatorial phosphorylation impacts tau’s behavior in both contexts is poorly understood. We hypothesized that contrasting developmental and pathological models of tau phosphorylation would yield insights into mechanisms of impaired neuronal resilience in tauopathies. We use transgenic PS19 tau mice to demonstrate that increases in individual tau phospho-epitopes with aging do not correlate with their enrichment in tau aggregates in these mice. Using seeded aggregation of tau in neurons, we observe that tau aggregate formation does not necessarily result in MT destabilization; instead, aggregation diminishes tau’s ability to counteract destabilization when MTs are externally perturbed. We identify that combinations of tau phospho-signatures are responsive to MT destabilization, and phosphorylation at residues S235 and S262 as particularly critical for regulation of interactions with MTs. Phosphorylation of these residues was associated both with significantly reduced binding to MTs, as well as impaired phosphatase response in aggregates. As a whole, this work provides mechanistic insight into phospho-regulation of tau function and prompts reconsideration of the canonical view that tau phosphorylation precedes MT destabilization in disease. Rather, we provide compelling evidence that phosphorylation of tau may be a response to changes in MT stability and that cell death may result from aggregated tau’s inability to function in this manner.

GRAPHICAL ABSTRACT
Schematic model of our findings: Tau phosphorylation is dynamically regulated in specific combinatorial patterns in response to shifting MT growth cycles.

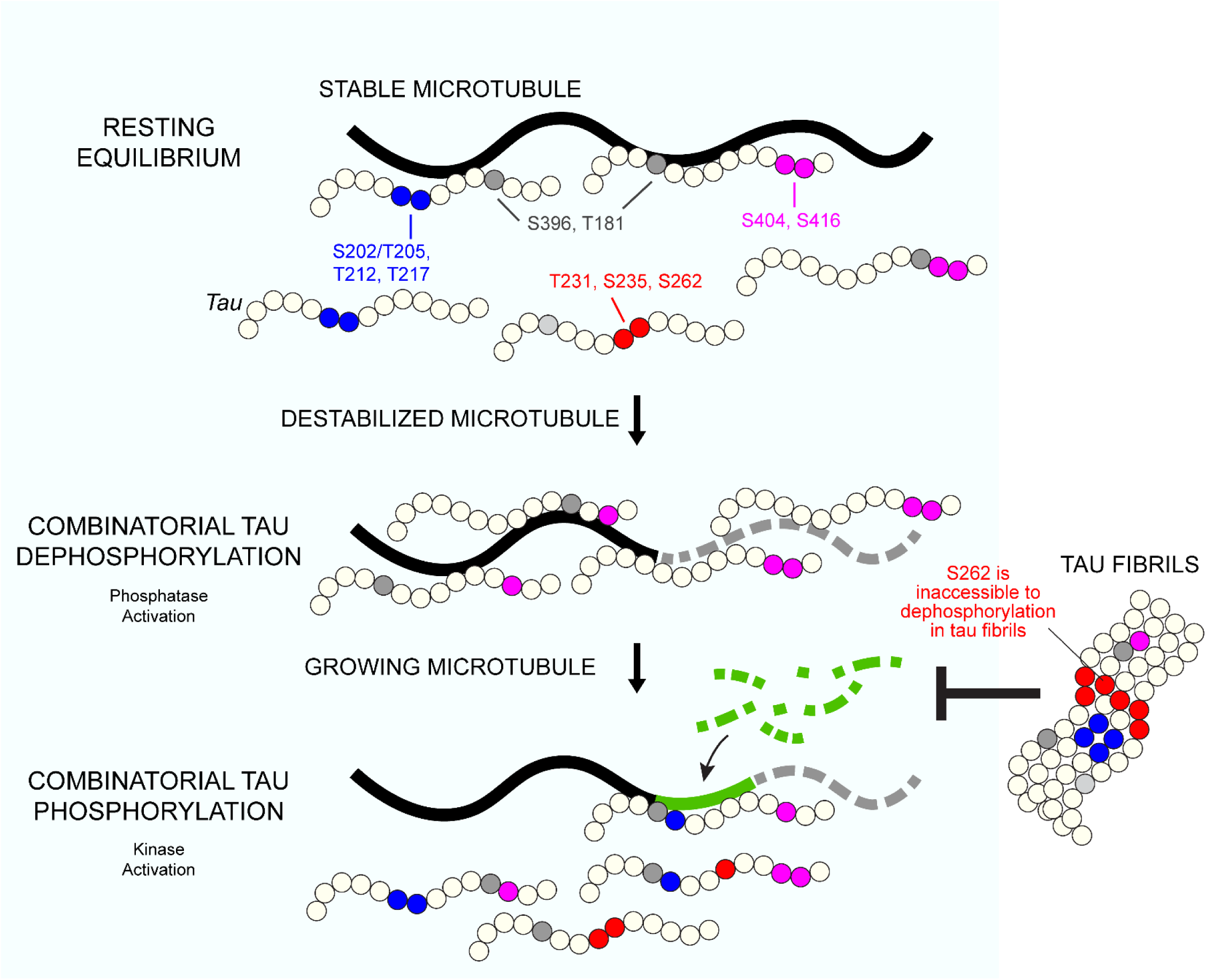

## INTRODUCTION

Hyper-phosphorylation of the microtubule (MT)-associated protein tau is strongly correlated with neuropathology in tauopathies including Alzheimer’s disease (AD)^1-3^. Substantial evidence supports that tau’s properties are significantly altered by post-translational modifications (PTMs) at specific residues, and that accumulation of these phosphate groups is found in intracellular deposits of aggregated tau. There is considerable discrepancy over whether any of these phospho-sites, of which there are over 80 potential sites in tau, are truly causative for aggregation^4^ or merely correlated with disease progression. However, several sites are canonically associated with fibrillar aggregates found in disease. These include phosphorylation at sites primarily throughout the proline-rich region (PRR; comprised of domains P1 and P2) and the C-terminal domain at residues T181, S202, T205, T212, S214, T217, T231, S235, S262, S396, S404, S416 and S422 (**Fig. 1A**). Phosphorylation at sites T181 and T217 are currently the most precise tau biomarkers for AD diagnosis^5-8^.

**Figure 1.**
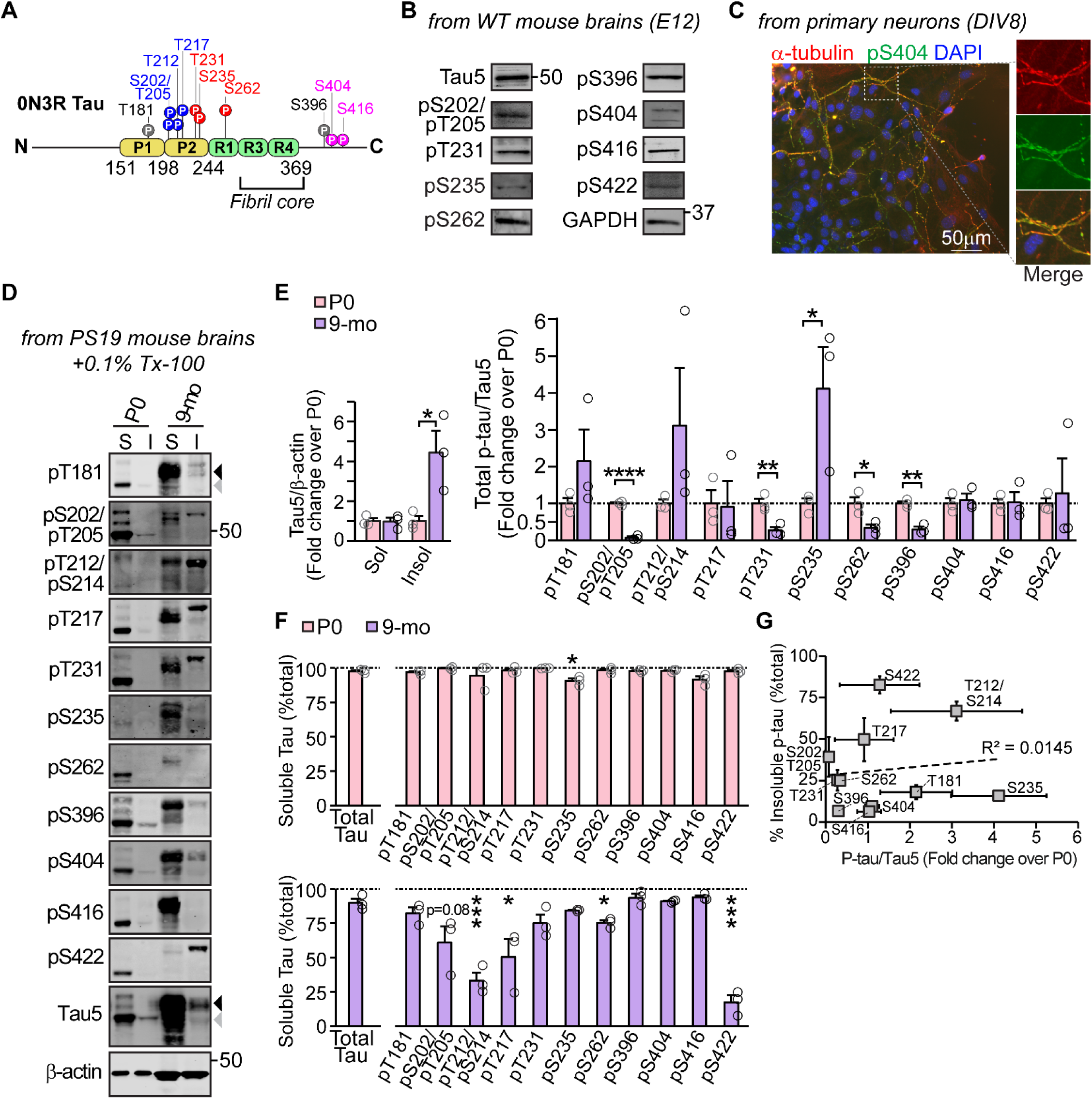
Tau phosphorylation levels in aged tauopathy mice do not correlate with its aggregation. **a)** Schematic of the fetal human 0N3R tau isoform. Residue numbering is based on the 2N4R isoform, as is convention in the field. The phospho-epitopes that were probed in this study in iNeurons are indicated and consistently color-coded throughout. **b)** Brains were collected from wildtype mice on embryonic day 12 (E12). Following separation by SDS-PAGE, pan-tau (anti-Tau5) and p-tau sites were detected by immunoblot at the indicated epitopes. Immunoblots are representative of 6 embryonic mouse brains. **c)** Tau phosphorylated at S404 (green) colocalized with MT protein α-tubulin (red) in wildtype primary mouse neurons at *DIV*8. For **d-g**, PS19 mouse brains were collected at post-natal days 0 (P0) and at 9 months. N=3 littermates per time point. The band shift between P0 and 9 months represents 0N3R mouse tau (gray arrowhead) and 1N4R human tau (black arrowhead). **d)** Triton-X solubility from PS19 mouse brains was used to separate [S]oluble from [I]nsoluble p-tau, followed by immunoblot. **e)** Pan-tau (anti-Tau5) solubility was quantified by Tau5/β-actin as a fold-change of P0 levels for both the soluble and insoluble fractions. Total p-tau levels (soluble + insoluble) were quantified and normalized to pan-tau (anti-Tau5) at each time point and represented as a fold-change of P0 levels. **f)** P-tau solubility was quantified as a percentage of total p-tau (soluble + insoluble) for each epitope and plotted alongside pan-tau solubility levels at each time point. Significance calculations were made compared to pan-tau solubility at each given age. **g)** The fold change in p-tau/Tau5 levels at 9-months compared to neonates (from ‘Fig. 1e’) was correlated with insoluble p-tau levels at 9-months of age (from ‘Fig. 1f’). Immunoblots and quantifications are representative of n=3 littermates. All data are representative of mean ± SEM. *P<0.05; **P<0.01; ***P<0.001; ****P<0.0001 by Student’s t-test.

The relationship between tau phosphorylation and its physiological state is complex. In numerous organisms, including humans and rodents, tau is highly phosphorylated during normal fetal brain development, underscoring a functional, regulatory role for many phosphorylation sites^9-12^. In fact, many of the same phospho-sites that are canonically associated with tau aggregates in disease are also present at this stage of development, when phospho-tau (p-tau) is highly soluble and functional. Adding to the complexity is that there are 6 tau isoforms expressed in the adult CNS. The shortest tau isoform, 0N3R (**Fig. 1A**), is exclusively expressed in fetal tissue, whereas isoforms 0N3R, 0N4R, 1N3R and 1N4R comprise the majority of tau in adult brains^13^. However, much of the biochemical tau research has focused on the longest isoform, 2N4R, despite its relatively low fraction in adult brains and negligible presence in fetal brains. The extent to which phospho-regulation of tau function is isoform-dependent is underexplored, but may be particularly relevant in the fetal brain, as the alternatively spliced regions are highly charged (net negative for N-terminal inserts, N; net positive for the repeat-regions insert, R), and thus their inclusion/exclusion may be integral to the effects of negative charges or other physico-chemical changes conferred by phosphorylation.

While the MT-binding domain (MBD) of tau has been the focus of intense research in promoting tau aggregate assembly, the adjacent flanking PRR and R’ domains are the primary regulatory regions of tau-MT dynamics. The PRR is the most abundantly phosphorylated domain in tau^14^. Compared to the MBD, the PRR is more competent in promoting MT assembly from soluble tubulin^15^, and tau binding to both soluble tubulin and MTs is greatly diminished by phospho-mimic mutations to the PRR^16,17^. The presence of the PRR is critical for tau’s ability to bind to MTs^18^; in fact, the MBD shows only weak affinity for MTs on its own^15,19^. Therefore, charge alterations to the PRR by phosphorylation are highly regulatory to tau-MT interactions, likely in a site-specific manner^20-22^.

Broadly, phosphorylation to tau is thought to enhance its dissociation from MTs, or inhibit its binding to growing MTs, to facilitate axonal outgrowth during development. Elevated levels of tau phosphorylation have been observed in fetal human brains^11,12^, supporting its physiological role in early neurodevelopment. Experimental strategies to increase tau phosphorylation – such as *in vitro* incubation of recombinant tau with kinases^23^ or phosphatase inhibition via okadaic acid treatment^24^ – can also induce high levels of tau phosphorylation in non-neuronal cell culture models. However, these methods are generally non-specific to phospho-epitopes in tau and may affect multiple substrates beyond tau. The complexity of tau phosphorylation is underscored by its regulation through both physiological and pathological processes. For instance, phosphorylation at one site may prime or enhance phosphorylation at additional sites, illustrating the coordinated action of multiple PTMs to a single tau molecule^25-28^. In disease contexts, phosphorylation at specific residues, such as S262 and S356, has been reported to exert opposing effects on tau’s assembly into aggregates^29-32^. Such conflicting findings may reflect the influence of combinatorial PTMs rather than the effects of individual phosphorylation events in isolation.

Furthermore, the mechanisms of compromised neuronal resilience downstream of tau aggregation remain incompletely understood. A prevailing hypothesis is that increased tau phosphorylation and its aggregation reduce tau-MT binding and thus result in destabilized MTs, thereby impairing neuronal health. This view is primarily based upon post-mortem studies reporting overall reductions in MT stability in AD brains^33^. However, closer scrutiny shows no detectable difference in MT stability between neurons containing paired helical filaments and those without^34^. Consistent with this finding, tau knockdown does not destabilize MTs in neuronal cultures; instead, tau appears to associate preferentially with the labile, rather than stable, MT fraction, potentially as a protective mechanism to buffer the MT network against stress^35,36^.

Here, we propose to revise the cascade of events in tau pathology by providing a mechanistic link between tau aggregation and alterations to MT stability in neurons. We show that fluctuations in MT growth cycles reshape tau’s phosphorylation landscape into specific combinatorial patterns that are canonically associated with insoluble tau. Contrary to longstanding assumptions, in our model system, tau aggregation does not result in destabilized MTs but instead disrupts the neuron’s ability to counteract MT destabilization. This loss-of-function is associated with specific phospho-epitopes, especially at S235 and S262, two modifications which significantly disrupt tau from binding to and stabilizing MTs. Our results point to the existence of distinct pools of tau defined by their combinatorial phospho-patterns. Overall, our findings suggest that combinatorial phosphorylation is not only a mechanism for regulating tau function in healthy neurons, but also a potential driver of tau’s loss-of-function in disease.

## RESULTS

Throughout this study, we used validated antibodies against specific p-tau sites^37-39^. We also validated these antibodies in mouse brains and human iNeurons by tau knockout and knockdown, respectively.

### Increases in single phospho-tau epitopes do not correlate with tau aggregation in mouse brains

Although phosphorylation states have been extensively characterized in tau aggregates, far less is known about how functional phosphorylation patterns are regulated in the developing brain. To begin addressing this gap, we first asked whether individual phosphorylation sites differ between developmental and disease states.

We began by verifying the presence of physiologically phosphorylated tau in fetal mouse brains, in confirmation of prior literature^10-12^. P-tau was readily detected as early as 12 days post-gestation (embryonic day 12, or *E12*) in wildtype embryos, with robust immunoreactivity at several canonical AD-relevant p-tau epitopes, including: S202/T205, T231, S235, S262, S396, S404, S416 and S422 (**Fig. 1B**). Tau phosphorylated at S404 was also detected in embryonic primary neuron cultures (**Fig. 1C**). Notably, the localization patterns for pS404 and α-tubulin were extremely similar, suggesting that tau that is phosphorylated at S404 remains capable of associating with MTs.

Because tauopathies such as AD are characterized by an aberrant, age-related increase in p-tau, we next compared tau phosphorylation patterns between neonatal mouse brains – representing a developmental stage with physiologically high levels of tau phosphorylation – and aged PS19 tauopathy mouse brains – a time when tau phosphorylation is largely ectopic. The PS19 tauopathy mouse model overexpresses the 1N4R isoform of tau harboring the pathogenic P301S mutation and develops insoluble tau deposits by 3 months of age^40^. We examined p-tau forms in neonatal (P0) and 9-month-old PS19 mouse brains by fractionating soluble and insoluble proteins and probing them using a panel of 11 site-specific p-tau antibodies (**Fig. 1D**).

Although most tau remained soluble at both times points (97.6% at P0; 89.7% at 9-months), the overall abundance of insoluble tau, as quantified by pan-tau (Tau5), was approximately 4-fold more in the 9-month-old compared to neonatal PS19 mouse brains (**Fig. 1D,E**). Numerous p-tau epitopes were detectable in aged PS19 mice, and a clear mass shift distinguished neonatal 0N3R tau from 1N4R tau at 9-months. The extent of phosphorylation was quantified by normalization to pan-tau to reveal heterogeneity across phospho-epitopes at 9-months relative to P0. To illustrate, phosphorylation at sites T212/S214 (AT100 epitope) and S235 increased markedly with age (>3-fold and >4-fold increase, respectively); in contrast, phosphorylation levels were a fraction of those found in neonates at sites S202/T205 (∼93% lower), T231 (∼72% lower), S262 (∼64% lower) and S396 (∼70% lower). No significant differences were observed between P0 and 9-months at sites T181, T217, S404, S416 or S422, suggesting comparable phosphate occupancy at these sites across both timepoints. Together, these results highlight conserved and divergent patterns of tau phosphorylation that may distinguish physiological from pathological modifications.

Next, we sought to investigate whether p-tau was preferentially enriched in insoluble tau, by comparing soluble p-tau against soluble total tau (**Fig. 1D,F**). In this quantification, we expected that in the absence of preferential enrichment, the fraction of p-tau found in the soluble fraction would be very similar to the total fraction of soluble tau protein. For the neonatal brains, we observed almost exactly this, with the fraction of soluble p-tau indistinguishable from soluble total tau across epitopes. However, at 9 months, we found several phospho-epitopes were more abundant in the insoluble fraction, especially at S202/T205 (∼61% soluble), T212/S214 (∼33% soluble), T217 (∼50% soluble) and strikingly at S422 (∼17% soluble). Additional phospho-sites such as T181 (∼82% soluble), T231 (∼75% soluble), S235 (∼84% soluble) and S262 (∼75% soluble) showed moderate enrichment in the insoluble fraction with age. In contrast, phosphorylation at C-terminal sites S396 (∼94% soluble), S404 (∼91% soluble) and S416 (∼94% soluble) largely reflected the fraction of soluble tau. In the 9-month-old mice, insoluble p-tau species were frequently mass-shifted upwards compared to soluble p-tau, consistent with more heavily modified insoluble tau species.

Collectively, these data indicate that although neonatal and tauopathy aged brains share many phospho-epitopes, fetal p-tau is generally not found in aggregates. This divergence between physiological phosphorylation in the fetal brain and ectopic phosphorylation during aging may give rise to distinct p-tau species, some of which may be unique to either state. However, changes in total p-tau abundance with age did not correlate with enrichment in the insoluble fraction (**Fig. 1G**). To illustrate, phosphorylation increased at T212/S214 by >3-fold and decreased at S202/T205 by >90% with age, yet both epitopes were strongly enriched in the insoluble fraction. These findings indicate that single phospho-epitopes cannot reliably predict tau aggregation, underscoring the need to evaluate combinatorial phosphorylation rather than individual sites in isolation.

### Human iNeurons provide a platform for contrasting physiological and pathological p-tau

The lack of clear, single-epitope distinctions between developmental and pathological tau phosphorylation *in vivo* prompted us to seek a neuronal culture model that would allow us to examine tau phosphorylation not only at individual sites, but also in combination. However, endogenous, non-overexpression models for studying tau phosphorylation have not been widely reported. Extensive phosphorylation to the fetal tau isoform, 0N3R, is reported in multiple organisms including developing avians^9^, rodents^10^ and humans^11,12^, suggesting that a naïve neuronal model may naturally support high levels of physiological p-tau. We therefore turned to human iPSC-derived induced neurons (iNeurons), which recapitulate key features of the embryonic environment^41-43^, for our studies.

Using iNeurons, we applied top-down mass spectrometry to quantify phosphorylation stoichiometries on intact endogenous tau. High levels of p-tau were readily detected, with a mean of 12.2 and median of 12.0 phosphates per tau protein. Phosphorylation levels were quantitatively characterized by four distinct clusters of 1-6, 7-10, 11-18, or 19-21 phosphates per tau molecule (**Fig. 2A & S1**). Within each cluster, the mode was 5, 8, 14 and 19 phosphates per tau. Acetylation was detected only on tau molecules modified with at least 11 phosphates (**Fig. S1**). Based on past qualifiers that >7 phosphates constitute tau hyperphosphorylation^12^, these data indicate that ∼82% of endogenous fetal tau in iNeurons meet this criterion. Thus, iNeurons express diverse, highly phosphorylated populations of tau molecules that are modified in potentially distinct combinatorial phospho-states.

**Figure 2.**
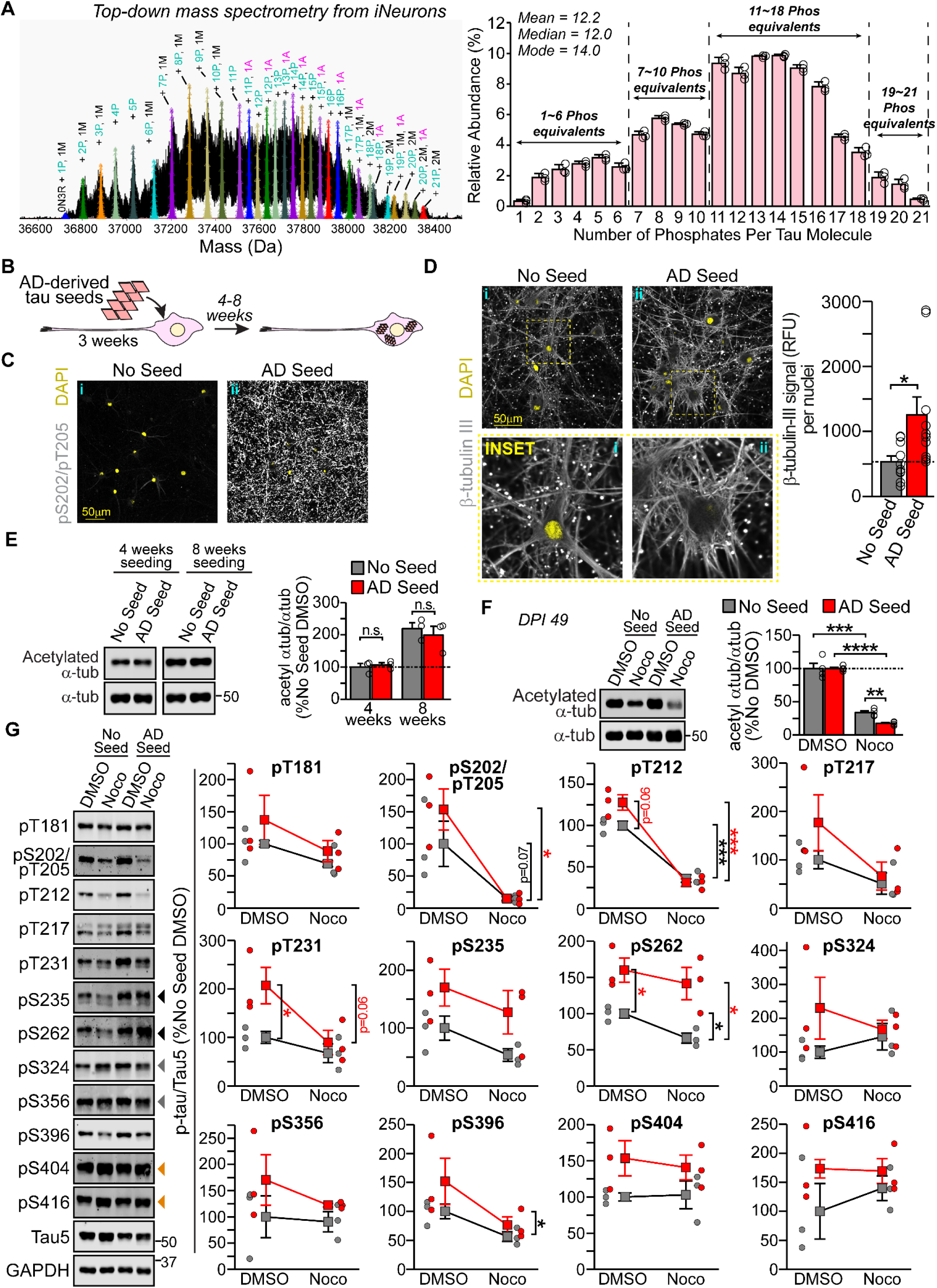
Tau aggregation disrupts the neuron’s ability to counteract MT destabilization. **a)** Tau was immunoprecipitated from iNeurons (*DPI 30*) with two pan-tau antibodies (Tau5 and HT7), and stoichiometries of phosphorylation events per tau protein were measured by top-down mass spectrometry with I^2^MS technology, indicating up to 21 phosphates per tau protein. PTMs were assigned based on mass shifts of ∼80 Da for phosphorylation, ∼42 Da for acetylation and ∼14 Da for methylation. The relative abundance of phosphate occupancy was calculated as a percentage of all detected 0N3R tau proteoforms. The annotated spectrum is representative of a single replicate of n=3 independent I^2^MS injections. Quantifications represent mean ± STD for n=3 independent I^2^MS injections. P = phosphorylation. A = acetylation. M = methylation. **b)** iNeurons were treated at 3 weeks post-differentiation with insoluble tau seeds derived from a post-mortem AD brain. Neurons were used for biochemistry or immunocytochemistry 4-8 weeks post-seeding. **For c-d**, soluble proteins were extracted from iNeurons by ice-cold methanol fixation at 6 weeks post-seeding (*DPI 63*). Images are representative of n=8 independent treatments. **c)** Immunocytochemistry was performed against anti-AT8 (pS202/pT205) at 40x magnification. Tau pathology was robust in neurons seeded with AD-derived tau. **d)** Immunocytochemistry was performed against anti-β-tubulin III at 60x magnification in neurons treated with either DMSO (control) or 10 µM nocodazole for 90 minutes. Quantification of β-tubulin III signal intensity from was performed using the mean signal intensity normalized to the number of nuclei in each field of view, from 10 random fields of view, for each sample group. **e)** Immunoblot and quantification of K40-acetylated α-tubulin normalized to α-tubulin 1A levels at 4-weeks (*DPI 49*) or 8-weeks (*DPI 77*) post-seeding. Blots and quantifications are representative of n=3 independent seeding and treatment experiments. For **f-g**, unseeded and seeded iNeurons were treated at 4 weeks post-seeding with either DMSO (control) or 10 µM nocodazole for 90 minutes, then harvested in RIPA buffer for Western blot. **f)** Immunoblot and quantification of K40-acetylated α-tubulin normalized to α-tubulin 1A levels. Blots and quantifications are representative of n=4 independent seeding and treatment experiments. **g)** Immunoblot against tau phosphorylated at the indicated phospho-epitopes. Blots and quantifications represent p-tau signal normalized to pan-tau for n=3 independent seeding and treatment experiments. The black arrowheads indicate the sites that responded uniquely to nocodazole treatment in seeded neurons. The gray arrowheads indicate sites that are contained with the tau fibril core. The orange arrowheads indicate two C-terminal sites that were differentially regulated in seeded versus unseeded neurons. All data are representative of mean ± SEM. *P<0.05; **P<0.01; ***P<0.001; ****P<0.0001 by Student’s t-test. N.S. = not significant.

Using a panel of tau phospho-epitope antibodies, as in ‘Figure 1’, we detected a number of the same AD-relevant phosphorylation modifications to tau in iNeurons as in brain tissue, including at sites T181 (AT270 epitope), S202/T205 (AT8 epitope), T212, T217, T231 (AT180 epitope), S235, S262, S396, S404 and S416 (**Fig. S2A**). Despite exclusively using previously validated p-tau antibodies, their specificity has not been confirmed in iNeurons. To address this, we generated a tau knockdown (tau^KD^) iPSC line using CRISPR-interference. Following neuronal differentiation, tau expression was nearly abolished relative to a non-targeting control (NTC) sgRNA (**Fig. S2B**), and unlike tau knockout models, tau knockdown was not compensated by upregulation of other MT-associated proteins (MAP1A, MAP1B, MAP2) (**Fig. S2C**). A further advantage of this tau^KD^ model is the ability to reintroduce tau by lentiviral transduction at approximately endogenous levels on a clean background (**Fig. S2D**), enabling measurements on individual isoforms without confounding compensation by other MAPs. Importantly, p-tau antibody specificity for tau was clearly demonstrated, apart from epitopes T212/S214, S214 (independently) and S422 (**Fig. S2E**).

Together, these results establish human iNeurons as a robust, endogenous model of fetal tau phosphorylation. The high stoichiometries of phospho-epitopes underscore the physiological importance of combinatorial phosphorylation during neuronal development and provide a powerful platform for contrasting physiological and pathological p-tau signatures.

### Tau aggregation impairs the neuron’s ability to counteract MT destabilization

To investigate how tau aggregation affects tau phosphorylation and MT stability, we next established a tau pathology model in human iNeurons by seeding endogenous tau using either AD brain-derived insoluble tau (**Fig. 2B**) or recombinant 3R tau seeds (**Fig. S3A**). Four to eight weeks after seeding, tau aggregation was robustly induced with AD brain-derived seeds (**Fig. 2C-ii & S3B-ii**) and less so with recombinant 0N3R seeds (**S3B-iii**), appearing as puncta or elongated structures within the soma and neurites, which were absent from unseeded controls (**Fig. 2C-i & S3B-i**).

Tau aggregation is thought to reduce MT stability by depleting soluble tau that would normally be associated with MTs. Thus, we expected to observe MT destabilization in our pathology model. Instead, we found the opposite: MTs remained largely intact in neurons harboring abundant tau aggregates (**Fig. 2D & S3C**). Following extraction of soluble tubulin, quantification of polymerized tubulin revealed a significant increase in total MT levels following seeding (**Fig. 2D & S3D**). Western blot against K40-acetylated α-tubulin, a marker of stable MTs, further corroborated that MTs remained stable as long as 8 weeks post-seeding (**Fig. 2E & S3E**).

This unexpected observation led us to revisit tau’s physiological function. Given some evidence that tau preferentially associates with labile tubulin and protects MTs from destabilizing enzymes^35,36,44^, we hypothesized that tau may serve not only to stabilize MTs but to buffer the MT lattice against destabilizing stress. If so, sequestration of tau into aggregates should reduce this protective capacity.

To test this, we challenged the MT network with nocodazole, a small molecule that destabilizes MTs by sequestering soluble tubulin heterodimers. Nocodazole disrupted MTs in all conditions, but the magnitude of MT loss was markedly greater in seeded neurons (**Fig. 2F** & **S3C,F**). Following nocodazole treatment, K40-acetylated α-tubulin decreased by 48% in AD-seeded neurons and 37% in 3R-seeded neurons compared to unseeded controls. As a whole, these data support a model in which tau aggregation renders neurons more vulnerable to MT destabilization, consistent with the loss of tau’s endogenous ability to counteract MT stress.

We next asked whether any specific p-tau epitopes could be connected to this phenotype. We quantified changes to site-specific tau phosphorylation levels in DMSO- and nocodazole-treated conditions, with and without seeding (**Fig. 2G & S3G**). In unseeded neurons, MT destabilization by nocodazole resulted in general trends toward dephosphorylation at many epitopes, consistent with MT destabilization triggering tau-targeted phosphatase activity to increase tau binding to soluble tubulin and/or MTs. In seeded neurons, the baseline levels of tau phosphorylation were higher at most epitopes relative to unseeded neurons. Remarkably, the seeded neurons exhibited a similar pattern as the unseeded ones following nocodazole treatment, with p-tau levels reduced to those of unseeded neurons, except at two epitopes, S235 and, most strikingly, S262. At S262, nocodazole induced a ∼31.5% reduction in phosphorylation in unseeded neurons but substantially less dephosphorylation in seeded neurons.

These results challenge the canonical tau cascade by demonstrating that increased tau phosphorylation and aggregation are not sufficient to significantly destabilize MTs. Rather, our data supports a model in which tau aggregation disrupts the neuron’s ability to buffer the MT network, increasing neuronal susceptibility to MT stress, likely primarily due to unavailability of sufficient amounts of soluble, functional tau. Dephosphorylation to increase MT affinity of tau may also be dysregulated at critical tau phospho-epitopes, especially S262.

### Phospho-sites are co-regulated in specific patterns in response to MT growth cycles

In general terms, phosphorylation to tau is thought to decrease binding of tau to MTs, facilitating MT depolymerization. In the developmental context, neurons maintain highly dynamic MTs, which undergo cycles of growth and shrinkage; our results here also show that they have high stoichiometries of tau phosphorylation (**Fig. 2A**). However, our seeding experiments suggest that tau phosphorylation levels can also be altered in response to changes in MT stability, not solely preceding MT destabilization.

We sought further insight into the response of p-tau changes to MT stability by pharmacologically perturbing the neuronal cytoskeleton and measuring the impact on site-specific tau phosphorylation. MTs were destabilized using either nocodazole or demecolcine or stabilized using paclitaxel; as a control, actin filaments were depolymerized using latrunculin A (**Fig. S4**). For clarity across all experiments, we consolidated phospho-sites responses into four categories based on observed trends by the following color scheme: gray (T181, S396), blue (S202/T205, T212, T217), red (T231, S235, S262) and magenta (S404, S416). We consistently maintained this consolidated grouping (also seen in **Fig. 1A**) throughout the remainder of the analyses. Data corresponding to each independent phospho-epitope are available in the supplementary information.

Consistent with our seeding results, MT disruption by demecolcine reproduced the effects of nocodazole by inducing pronounced tau dephosphorylation at numerous phospho-epitopes including S202/T205, T212 and T217, and more modest dephosphorylation at T231, S235 and S262. Intriguingly, MT disruption resulted in increased tau phosphorylation in the C-terminus at epitopes S404 and S416. MT stabilization by paclitaxel produced similar tau dephosphorylation patterns at epitopes S202/T205, T212, T217, though effects at T231, S235 and particularly S262 were less pronounced. In contrast, actin depolymerization had minimal effects on tau phosphorylation at most of the measured epitopes, indicating that these signatures are primarily driven by MT perturbation. Together, these results suggest that tau phosphorylation is differentially regulated in response to changes to MT stability, rather than acting solely as its upstream effector.

We further expanded upon this finding to measure how tau phosphorylation is modulated in response to MT collapse and re-stabilization. In this paradigm, acute MT catastrophe was induced in iNeurons using a brief (5 minute) cold exposure followed by either short- or long-term nocodazole treatment for 15 or 90 minutes, respectively, to analyze the cellular response to acute versus sustained MT damage. After 90 minutes, nocodazole was washed out to allow MT recovery (**Fig. S5A**). We tracked site-specific phosphorylation to tau throughout the treatment and recovery periods.

As expected, nocodazole disrupted the spindle-like MT lattice and tau’s axonal localization, both of which recovered within 3 hours after nocodazole washout (**Fig. S5B**). These visual findings were corroborated by immunoblot against K40-acetylated α-tubulin and detyrosinated α-tubulin, an orthogonal marker of stable MTs, which were reduced to ∼50% of control levels and recovered to ∼75% at 3 hours after nocodazole washout (**Fig. 3A,B**).

**Figure 3:**
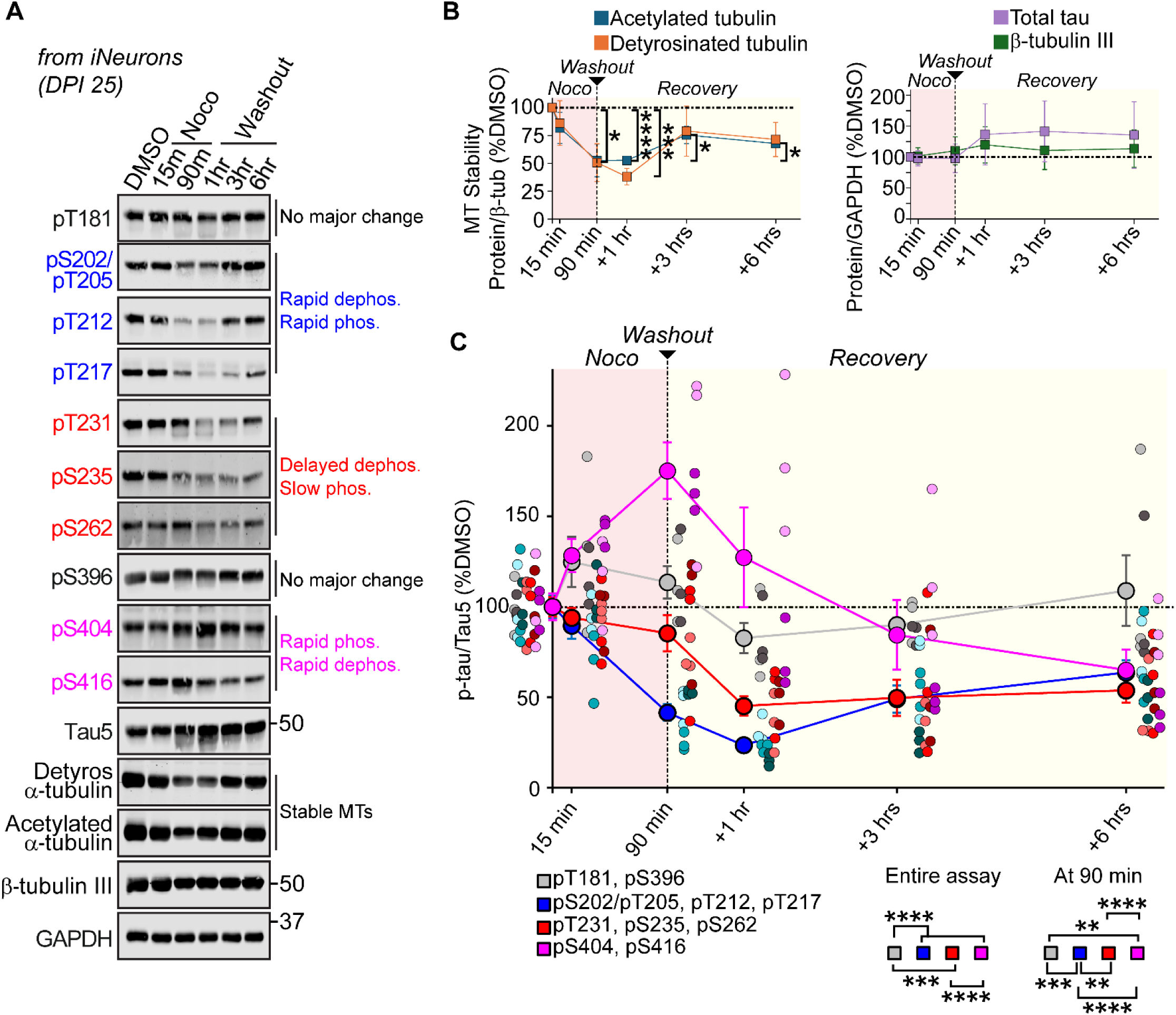
Phospho-sites throughout tau are differentially regulated to counteract MT destabilization. iNeurons (*DPI 25*) were treated with either DMSO (control) or 10 µM nocodazole for either 15 min, 90 min, or 90 min followed by nocodazole washout and recovery for 1 hour, 3 hours or 6 hours. **a)** Samples were harvested for SDS-PAGE followed by immunoblot against the indicated proteins. N=3 independent cultures and treatments per group. **b)** Quantification of ‘a’ for MT stability as measured by changes to detyrosinated α-tubulin and K40 acetylated α-tubulin during nocodazole treatment and washout. Total (pan) tau and β-tubulin III levels were not significantly impacted throughout the assay. *P<0.05; ***P<0.001; ****P<0.0001 by Student’s t-test. **c)** Quantification of ‘a’ for p-tau normalized to pan-tau. **P<0.01; ***P<0.001; ****P<0.0001 by post-hoc Tukey’s HSD test.

We then correlated dynamic changes in tau phosphorylation with MT collapse and re-stabilization (**Fig. 3A,C** and **S5C**). Tau phosphorylation dynamics could be classified into four distinct groups.

The first group, comprised of T181 and S396 (gray line), was not significantly impacted by nocodazole treatment or washout. The second group (S202/T205, T212, T217; blue line) consisted of rapid responders, which underwent strong dephosphorylation (∼23% of control levels) in response to nocodazole treatment followed by rapid re-phosphorylation after nocodazole washout (recovery to ∼64% of control levels). These epitopes closely mirrored the changes to MT stability throughout the time course. The third group (T231, S235 and S262; red line) consisted of delayed responders, which exhibited slower and more modest dephosphorylation in response to nocodazole treatment (∼45% of control levels), with limited levels of re-phosphorylation after nocodazole washout (recovery to ∼54% of control levels). These sites changed more slowly than the blue cluster, and continued to decrease until 1 hour into the recovery phase, even after detyrosinated α-tubulin and acetylated α-tubulin levels had stabilized. The fourth group (S404, S416; magenta line) consisted of inverse responders, which were rapidly phosphorylated within 15 min of MT collapse: phosphorylation at these sites continued to increase until the nocodazole was washed out (∼175% of control levels), at which point these sites were immediately, rapidly dephosphorylated (∼65% of control levels) in alignment with the timescale of MT stabilization, highlighting an inverse relationship with MT integrity. The prompt increase in phosphorylation at these two epitopes may reflect the importance of regulation at these sites for counteracting MT destabilization.

Nocodazole treatment did not impact total (pan) tau or β-tubulin III levels, though both proteins showed modest upregulation after nocodazole washout, suggesting a potential compensatory cellular response to counteract MT destabilization (**Fig. 3A,B**). Notably, changes to tau phosphorylation preceded any detectable increases in protein abundance; therefore, the observed patterns of dephosphorylation and phosphorylation were independent of new protein synthesis.

Overall, our data show that tau phosphorylation is dynamically and combinatorially regulated by MT stability. Co-regulated clusters, including T181-S396, S202/T205-T212-T217, T231-S235-S262 and S404-S416, may reflect distinct roles in MT collapse and recovery, supporting a model in which tau phosphorylation signatures act as responders to MT integrity rather than solely as drivers of MT destabilization.

### Tau dephosphorylation by PP2a is necessary to counteract MT destabilization

The co-regulated phosphorylation patterns identified above suggest that p-tau exists in distinct pools that respond differentially to MT stress. As MTs undergo collapse, many of the p-tau sites we probed are dephosphorylated, shifting the equilibrium toward increased tau binding to MTs as a compensatory mechanism to counteract MT catastrophe. As cellular conditions become conducive to MT recovery or repair, these same sites are re-phosphorylated, tipping the balance back toward less MT-bound tau and restoring dynamic flexibility. This regulatory model contrasts with the prevailing view in neurodegeneration that phosphorylation to tau primarily drives MT destabilization by disrupting tau’s MT-binding capacity. In contrast, our seeding experiments showed that even large increases in tau phosphorylation, accompanied by sequestration of tau into aggregates, did not result in destabilized MTs (**Fig. 2E**), raising the possibility that changes in MT stability precede and influence site-specific tau phosphorylation. Taken together, these observations suggest that tau phosphorylation can function both as a consequence and a driver of MT stability, with the directionality of this relationship potentially differing between physiological and pathological states.

To further investigate, we probed the molecular pathways that regulate tau dephosphorylation and phosphorylation during MT collapse and recovery. We first characterized the activation states of a phosphatase and kinases that have been previously implicated in tau regulation^10,25^. The phosphatase PP2a was activated during nocodazole treatment, coinciding with the period of tau dephosphorylation (**Fig. 4A,B**, *top panels*). The kinases Erk1/2 and Akt1 were activated after nocodazole washout, temporally aligning with the period of tau re-phosphorylation (**Fig. 4A,B**, *middle two panels*), whereas the kinase GSK3β was activated as soon as 15 minutes after nocodazole treatment (**Fig. 4A,B** *bottom panels*), preceding the period of phosphorylation of most tau sites except for S404 and S416.

**Figure 4.**
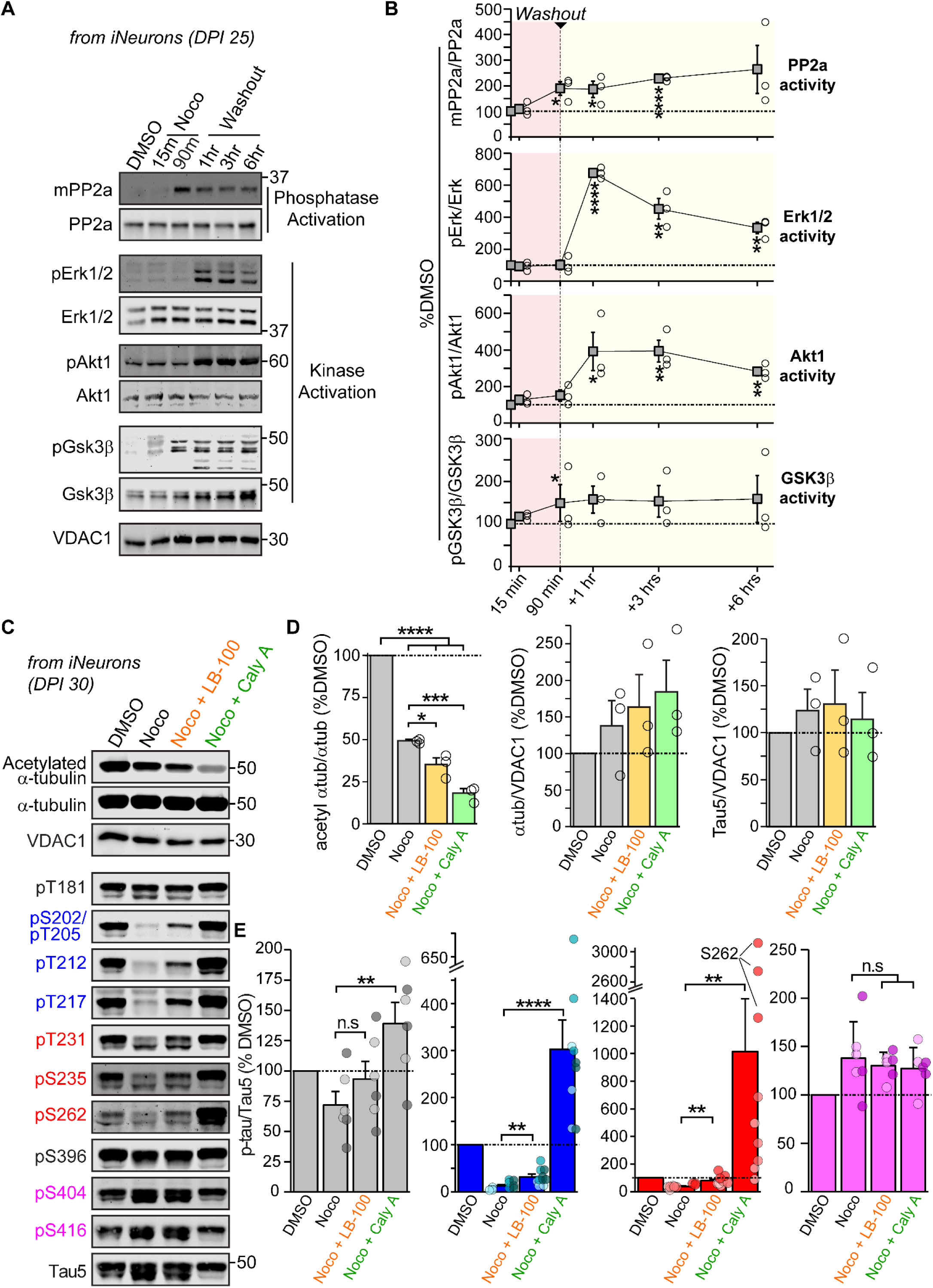
Tau dephosphorylation by PP2a is necessary to counteract MT destabilization. **a)** iNeurons (*DPI 25*) were treated with either DMSO (control) or 10 µM nocodazole for either 15 min, 90 min, or 90 min followed by nocodazole washout and recovery for 1 hour, 3 hours or 6 hours. PP2a activation was measured by methylation at Leu309. Kinase activation was measured by phosphorylation against Erk1/2 (T202/T204), Akt1 (S473) and GSK3β (Y216). **b)** Quantification of ‘a’. Markers for kinase and phosphatase activation were normalized to their respective pan-proteins: pErk1/2 (T202/T204) normalized to pan-Erk1/2, pAkt1 (S473) normalized to pan-Akt1, pGSK3β (Y216) normalized to pan-GSK3β, and mPP2a (L309) normalized to pan-PP2a. Blots are representative of n=3 independent treatments. Significance calculations were made compared to the initial timepoint. For **c-e**, iNeurons (*DPI 30*) were treated with either DMSO (control), 10 µM nocodazole, or 10 µM nocodazole supplemented with either LB-100 (20 µM) or Calyculin A (20 nM) for 90 min. N=3 independent cultures and treatments. **c)** Representative immunoblots are shown for the indicated proteins and p-tau epitopes. **d)** Quantification of changes to MT stability by immunoblot against K40-acetylated α-tubulin normalized to α-tubulin levels. No significant changes were detected to total α-tubulin or pan-tau (anti-Tau5) levels, normalized to VDAC1. **e)** As in ‘d’, changes to p-tau were measured by immunoblot against the indicated phospho-epitopes. P-tau levels were normalized to pan-tau (anti-Tau5) for each sample. N.S.=not significant. *P<0.05; **P<0.01; ***P<0.001; ****P<0.0001 by Student’s t-test. All data are representative of mean ± SEM.

We next asked whether inhibiting these enzymes could alter MT stability. We combined phosphatase inhibition with nocodazole treatment to assess the role of tau dephosphorylation during MT collapse, and applied kinase inhibition during the recovery period to test for the impact of blocking tau phosphorylation during MT recovery. Among the kinase inhibitors that we tested – targeting Akt1 (AKT1 inhibitor VIII), Erk1/2 (Erk1/2 inhibitor I) and GSK3β (BIO) – only the Akt1 inhibitor was both effective and selective for its intended target in iNeurons during the tested timeline (**Fig. S6A,B**). Akt1 inhibition resulted in only a modest improvement in MT stability compared to DMSO treatment (**Fig. S6B**). These results suggest that blocking tau phosphorylation at sites that are targeted by Akt1 during MT recovery has a limited effect on MT stabilization. Instead, these findings imply that the critical regulatory response to counteracting MT destabilization may not occur during re-phosphorylation but rather during the dephosphorylation phase, when tau must rapidly respond to MT damage.

Accordingly, we inhibited PP2a activity using either LB-100 (reversible) or Calyculin A (potent, irreversible) during nocodazole treatment. Both inhibitors effectively reduced PP2a activity (**Fig. S6C**). LB-100 treatment impaired MT stability during nocodazole treatment but MTs were able to recover after washout (**Fig. 4C,D & Fig. S6C**). In contrast, Calyculin A treatment led to severe MT destabilization, comparable to AD-seeded conditions (**Fig. 2F**), and MTs failed to recover after washout, consistent with the irreversible nature of inhibition.

Finally, we correlated changes in MT stability with epitope-specific changes in tau phosphorylation. Mild PP2a inhibition by LB-100 was compared to potent inhibition by Calyculin A to tease out specific tau phospho-epitopes that may be critical for MT recovery. PP2a inhibition correlated with blunted dephosphorylation at many epitopes (S202/T205, T212, T217, T231, S235 and S262), while sites T181, S396, S404 and S416 remained largely unaffected (**Fig. 4C,E & S7**). Of note, Calyculin A treatment resulted in especially higher levels of phosphorylation at sites T212, S235, and most prominently at S262. While we do not ascribe causal roles to individual epitopes, the disproportionate sensitivity of S235 and S262 to phosphatase inhibition, as well as their delayed re-phosphorylation following nocodazole treatment (**Fig. 3C & S5**), underscores that dephosphorylation of these residues was closely linked to the cell’s ability to restore MT stability following collapse.

Together, these findings demonstrate that PP2a activation is required for neurons to buffer against MT destabilization, and that distinct subgroups of tau phospho-sites are tightly linked to this response. This establishes that tau dephosphorylation is not passively correlated with counteracting MT destabilization but is a component of the neuronal response to MT catastrophe.

### Tau phosphorylation at T231, S235 and S262 directly and substantially disrupt tau-MT interactions

Our data demonstrate that fetal tau is modified at many of the same phospho-epitopes as pathology-associated tau, and in some cases to a greater extent, yet tau remains functional and MTs are stable in the developing brain. Given that different p-tau species were observed to be co-regulated in certain clusters in response to chemical destabilization of MTs, we asked whether these same clusters differ in their physical associations with MTs. Tau phosphorylation regulates tau-MT interactions, although molecular details are lacking for many individual phosphorylation sites, and combinatorial effects are even more poorly characterized. Therefore, we fractionated intact MTs from tau^KD^ iNeurons expressing 0N3R tau and compared phospho-site associations with MTs for tau.

Effective MT fractionation was indicated by separation of detyrosinated α-tubulin from neuron-specific enolase (NSE), a marker for neuronal cytosol and plasma membrane (**Fig. 5A & S8**). Approximately 77% of β-tubulin III and 43% of tau was found in the intact MT fraction.

**Figure 5:**
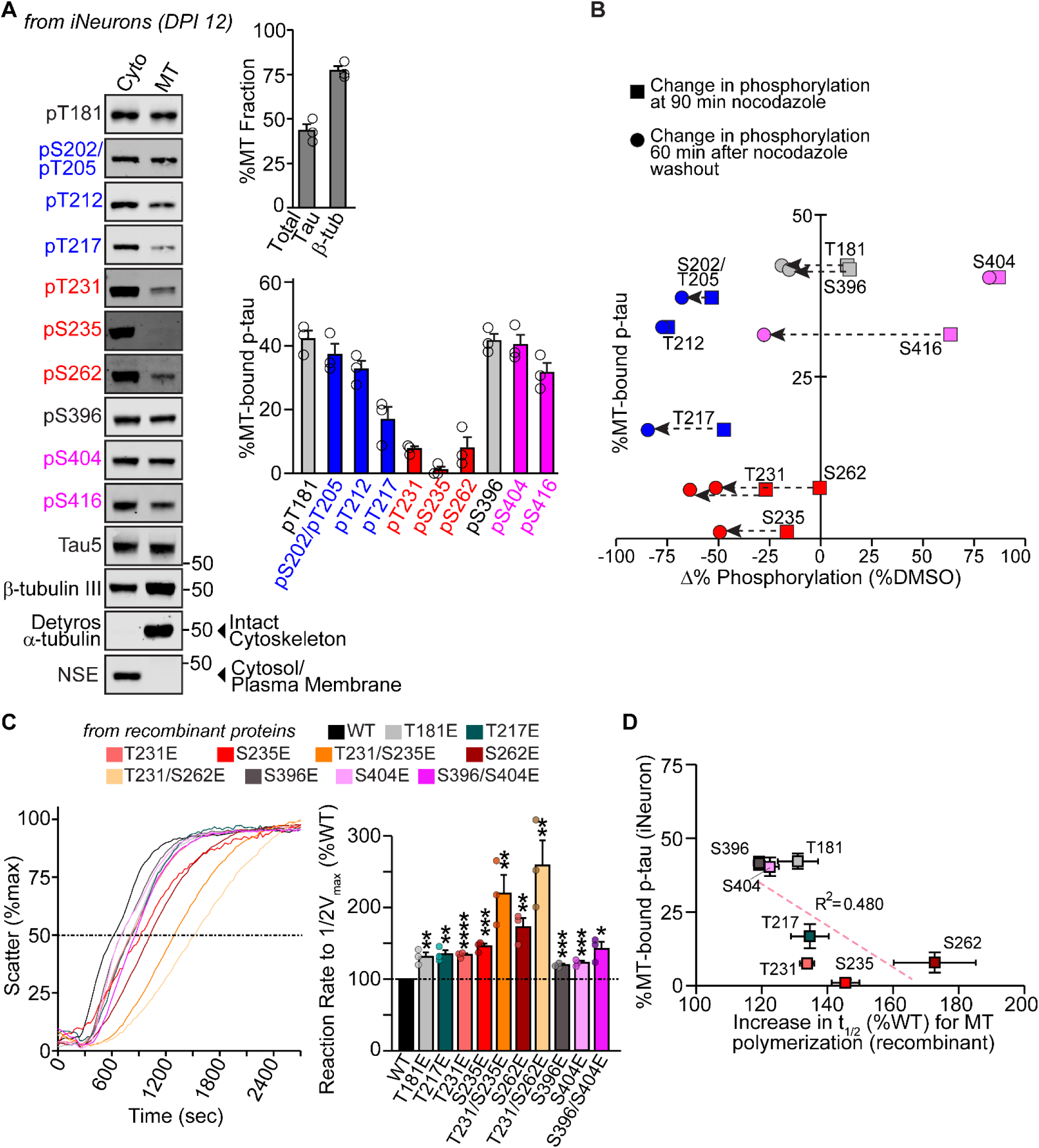
Site-specific phosphorylation directly disrupts tau’s interaction with MTs. **a)** MTs were fractionated from tau^KD^ iNeurons expressing 0N3R^myc^ at *DPI 12* (n=3 independent cultures, transductions and fractionations). SDS-PAGE followed by immunoblot was used to detect p-tau levels in cytosolic and MT-bound fractions. Separation of detyrosinated α-tubulin (MT fraction, only) from neuron-specific enolase (NSE), a cytosol-specific protein (cytosolic fraction, only), indicated successful separation of intact MTs from the cytosol. The amounts of pan-tau (anti-Tau5), β-tubulin III and p-tau that were found in the MT-bound fraction were quantified as a percentage of total protein (cytosolic + MT fractions) for each indicated target. **b)** The percentage of each p-tau epitope bound to MTs (from ‘Fig. 5A’) was plotted alongside the changes in phosphorylation relative to DMSO treatment at 90 minutes of nocodazole treatment and 1 hour after washout (from ‘Fig. 3D’), revealing the trends in dephosphorylation over time. **c)** MT polymerization was detected *in vitro* by light scatter at 340 nm for recombinant wildtype tau or phospho-mimics of each at the indicated sites. The reaction mixture (20 µM soluble tubulin + 10 µM recombinant tau in BRB80 buffer) was supplemented with 1 mM DTT (final), initiated by addition of 1 mM GTP (final) at 37C and monitored for 45 minutes on a spectrophotometer. N=3 independent polymerization reactions for each tau variant. Quantification of the change in the reaction rate to 50% V_max_ for each phospho-mimic was compared to wildtype tau. **d)** The percentage of MT-bound p-tau from iNeurons (from ‘Fig. 5a’) was plotted alongside the increase in t_1/2_ rate of MT polymerization by tau phospho-mimics compared to WT (from ‘Fig. 5c’), for epitopes T181, T217, T231, S235, S262, S396 and S404. All data are representative of mean ± SEM. *P<0.05; **P<0.01; ***P<0.001; ****P<0.0001 by Student’s t-test.

We then quantified the distribution of phospho-sites across both fractions (**Fig. 5A**). Tau phosphorylated at sites in the late P2 domain (T231, S235) and in the R1 domain (S262) was almost exclusively in the cytosol, indicating that phosphorylation at these sites is strongly associated with MT dissociation. In contrast, phosphorylation at more N-terminal P2 domain sites (T181, S202/T205, T212 and T217) and within the C-terminal domain (S396, S404, S416) was more strongly associated with MTs. These localizations are consistent with the idea that the PRR can regulate tau-MT binding independently of the MBD. Overall, we observed a bimodal distribution, in which tau that was phosphorylated outside of the last 15 residues of the P2 domain in the PRR + early MBD residues (T231, S235, S262) was more likely to be associated with the MT fraction compared to those within this region.

We then integrated these observations with our MT growth-cycle experiments to provide insight into the relationship between MT-association and responsiveness of tau to MT integrity (**Fig. 5B**). The cluster of sites that showed the slowest re-phosphorylation after nocodazole washout (T231, S235, S262) also exhibited the lowest levels of MT-binding. This alignment suggests that these sites must remain dephosphorylated for tau to effectively engage MTs during recovery. In contrast, epitopes that showed minimal changes or rapid re-phosphorylation after nocodazole washout (T181, S396, S202/T205, T212 and T217) were comparatively enriched in the MT-bound fraction, consistent with a model in which their phosphorylation state is less restrictive for MT association.

The alignment between the clusters of behavior of phospho-sites within these data sets reveals a hierarchy of which sites are most critical for regulating functional interactions of tau with tubulin/MTs. The striking sensitivity of S235 and S262 to both phosphatase inhibition and MT-binding assays reinforces their central position in regulating tau-MT interactions.

Thus far, these data demonstrate that tau phosphorylation changes are strongly correlated with its association with MTs and the observed alterations to MT stability in the presence of MT stressors. To directly test for a link between site-specific tau phosphorylation and MT stability, we turned to a cell-free system to model and validate these biophysical interactions.

While phospho-mimetic substitutions are widely used, they do not reproduce endogenous combinations of modifications, and their ability to accurately mimic bonafide phosphorylation remains uncertain. To address this, we cloned, expressed and purified wildtype 0N3R tau along with single-site phospho-mimetic substitutions (Thr◊Glu or Ser◊Glu) at key sites identified from our neuronal dataset: T181, T217, T231, S235, S262, S396 and S404. We also generated three combinatorial phospho-mimetics: T231E/S235E and T231E/S262E, as well as S396E/S404E based on its relevance to tau fibrils.

Using tubulin purified from bovine brains, we assessed the capacity of each tau variant to promote polymerization of soluble tubulin (**Fig. 5C**). Most single-site phospho-mimetics impaired MT polymerization compared to their wildtype counterparts. Of the individually tested phospho-mimetics, the S235E and S262E mutations of MT polymerization (∼46% and ∼73% reduction, respectively). Combinatorial phospho-mimetics revealed further that dual glutamate substitutions at T231/S235 and T231/S262 resulted in additive inhibition of MT polymerization activity: In fact, these combinations impaired tau activity more than would be predicted from the individual mutations. Consistent with our observations in neurons, phospho-mimetic mutations at S396E, S404E and in combination had comparatively modest effects on MT polymerization.

These experiments reinforce our finding that phosphorylation particularly at S235, S262 and in combination with T231 directly impair tau’s ability to bind to tubulin/MTs. In support, correlation of p-tau bound to MTs (from iNeurons) with MT polymerization rates (from recombinant proteins) emphasized a clear disruption by site-specific phosphorylation/phospho-mimics to tau’s (R^2^=0.480) interaction with MTs (**Fig. 5D**). Our data support prior predictions that S235 and S262 make more contacts with MTs relative to other phospho-epitopes, suggesting that phosphorylation at these residues plays a particularly critical role in modulating MT stability^20^.

### Tau is phosphorylated in site-specific combinatorial patterns in iNeurons

Finally, we asked whether the regulation of site-specific tau phosphorylation in iNeurons aligns with patterns of co-phosphorylation. In pursuit, we next asked whether phosphorylation patterns are random or occur in distinct combinations. This question is essential because Western blot-based measurements cannot resolve combinatorial PTMs on individual tau molecules. Meanwhile, our earlier measurements indicate that tau-MT interactions may be dependent upon combinatorial phosphorylation to tau rather than single modifications. Although several dual-phosphorylation events have previously been observed in tau aggregates (e.g., S202/T205, T212/S214, T231/S235 and S396/S404^45-49^), combinatorial occurrences of tau phosphorylation are generally understudied, in particular as relevant to tau function.

We began to quantify these patterns by first immunoprecipitating p-tau from tau^KD^ iNeurons expressing 0N3R^EGFP^ and back-calculating phosphorylation frequencies by comparing the recovery of phospho-specific tau with total tau captured in each immunoprecipitation^50^. This allowed us to estimate the proportion of tau molecules phosphorylated at a given epitope. In agreement with the top-down mass spectrometry results, we found abundant and heterogeneous phosphorylation levels: pT217 and pS235 were uncommon (<20%), pT181 and pT212 were moderately common (>20%), and pS202/pT205, pT231 and pS396 were abundant (>45%) (**Fig. S10**).

To investigate co-occurrence between phosphorylation sites, we asked: when tau is phosphorylated at a given site, what is the likelihood that it is also phosphorylated at a second site? To address this question, we immunoprecipitated p-tau at individual phospho-epitopes targeting prominent AD-relevant epitopes within the PRR (T181, S202/T205, T212, T217 and T231) and one epitope in the C-terminus (S396), then probed for co-occurring phosphorylation at additional sites (**Fig. 6A,B**). Co-occurrence frequencies were calculated as the signal for each secondary phospho-site (normalized to input) divided by the signal for the primary immunoprecipitated site (also normalized to input).

**Figure 6:**
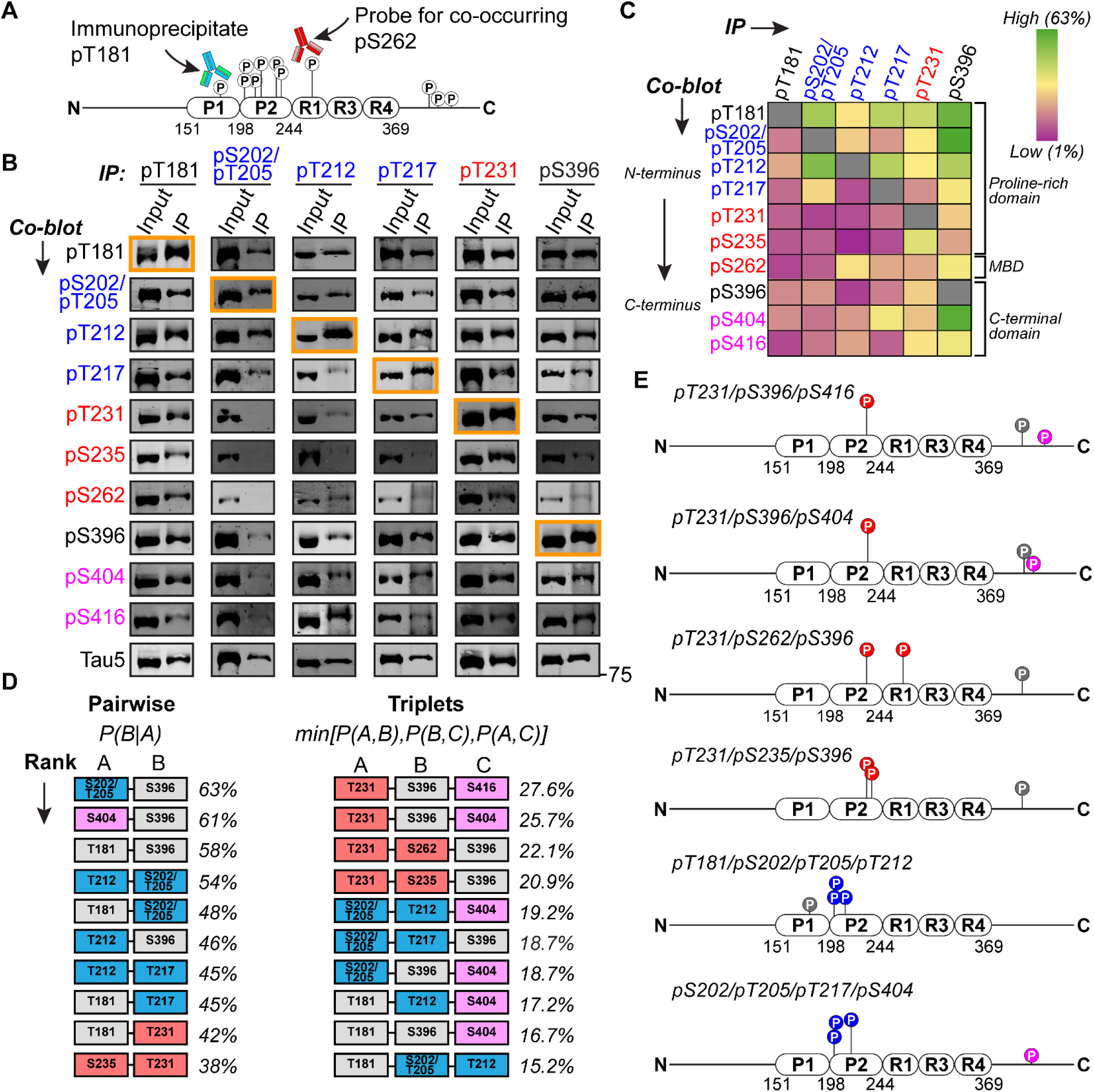
Phospho-epitopes are co-phosphorylated in specific patterns in iNeurons. **a)** General workflow for p-tau immunoprecipitation from iNeurons under denaturing conditions, followed by blotting for co-occurring phospho-epitopes. **b)** Tau^KD^ iNeurons were lentivirally transduced with 0N3R^EGFP^. P-tau was immunoprecipitated at 32 days post-induction, followed by separation by SDS-PAGE and immunoblotted against the indicated immunoprecipitated p-tau epitope and co-blotted at various p-tau epitopes. The immunoblot outlined in orange indicates the immunoprecipitated p-tau species in that given column of immunoblots. Blots are representative of n=4 (pT181, pT212, pS396), n=5 (pS202/pT205) and n=6 (pT217, pT231) independent cultures and immunoprecipitations. **c)** Heat map representing the quantification of phospho-site co-occurrence for each immunoprecipitated p-tau species, as described in the ‘Methods’ section. Green=high co-occurrence. Purple=low co-occurrence. Data is representative of n=4 (pT181, pT212, pS396), n=5 (pS202/pT205) and n=6 (pT217, pT231) independent cultures and immunoprecipitations. **d)** The top 10 most common combinations of two and three co-occurring phospho-epitopes are ranked from top to bottom. For pairwise combinations, the percentages adjacent to each combination reflect the highest linkage for a given pair of phospho-epitopes. Combinations of three phospho-epitopes were predicted by minimum linkage analysis as described in the ‘Methods’ section. The percentages adjacent to each triplet represent the lowest co-occurrence probability between any of the three indicated epitopes. **e)** Schematic representation of several predicted combinatorial phospho-modifications to 0N3R tau in iNeurons.

This approach yielded numerous unique patterns of co-occurring phospho-sites, indicating that tau phosphorylation follows non-random, site-specific patterns (**Fig. 6C**). Two previously established combinations, pS396/pS404 and pT231/pS235, were among the most frequently observed combinations, validating our approach. Specifically, phosphorylation at S396 conferred a ∼61% likelihood of concurrent phosphorylation at S404. Similarly, tau phosphorylated at T231 exhibited a ∼41% likelihood of co-occurring phosphorylation at S235.

We next sought to identify novel patterns of tau phosphorylation by ranking the top ten most common pairwise co-occurrences (**Fig. 6D**, *left column*). From highest to lowest co-occurrence, the most prominent dual combinations of phospho-epitopes were as follows (with the immunoprecipitated site <u>underlined</u>): pS202/pT205/<u>pS396</u> (68%); <u>pS396</u>/pS404 (61%); <u>pS202/pT205</u>/pT212 (61%); pT181/<u>pS396</u> (58%); pT212/<u>pT217</u> (54%); pT181/<u>pS202/pT205</u> (52%); pT212/<u>pS396</u> (46%); pT181/<u>pT217</u> (45%); pT181/<u>pT231</u> (42%); <u>pT231</u>/pS235 (41%). Additionally, tau phosphorylated at S262 exhibited 30% linkage with pT212.

Asymmetric linkages were also evident. For instance, pT217 was also phosphorylated at T212 in ∼54% of cases, but the reverse relationship was not as prevalent: only ∼8% of tau phosphorylated at T212 also carried phosphorylation at T217. Such asymmetric patterns may reveal potential priming or hierarchical ordering of tau phosphorylation events.

Next, we explored this dataset to infer potential combinations of three simultaneous co-occurrences. Pairwise probabilities cannot infer higher-order combinations; therefore, we applied minimum-linkage analysis to estimate the likelihood of triplet co-occurrences by selecting the weakest link among the triplets. Using this relatively conservative metric, the top ten most likely sets of three simultaneously co-occurring tau phospho-epitopes were ranked from highest to lowest (**Fig. 6D**, *right column*). Some of the most frequent co-occurrences, such as T231-S262-S396, may represent specific tau phospho-proteoforms (**Fig. 6E**).

The pairwise and triplet analyses revealed robust linkages between specific phospho-epitopes, aligning with the sites that we had earlier grouped into clusters. Frequent co-occurrences were observed between S202/T205, T212 and T217, forming a tightly associated cluster (color-coded blue). A second cluster emerged linking T231 with S235 and S262 (color-coded red). In contrast, phosphorylation at T181 and S396 was broadly distributed, indicating more promiscuous co-occurrence patterns (color-coded gray). Finally, S404 and S416 co-occurred most strongly with S396, and to a lesser extent with T217 and T212, forming a distinct subgroup (color-coded magenta).

These structured patterns suggest that tau phosphorylation is a coordinated process. The frequent co-occurrence of certain phospho-epitopes may reflect coordinated regulation by shared kinases or shared functional consequences in modulating tau’s biochemical properties and behavior. Accordingly, specific combinations of phosphorylation may define distinct pools of tau, each potentially associated with unique functional roles.

Importantly, the most prominent phosphorylation co-occurrences mirrored the patterns of dephosphorylation and re-phosphorylation in response to MT depolymerization and recovery (from **Fig. 3C**). For instance, pS202/pT205 and pT212 (∼61% co-occurrence), pT212 and pT217 (∼54% co-occurrence), pT231 and pS235 (∼41% co-occurrence), and pT181 and pS396 (∼58% co-occurrence) were similarly impacted by nocodazole treatment and washout. These coordinated changes support the notion that these sites are not only frequently co-phosphorylated but also co-regulated in response to shifts in MT stability. Combined with our functional data, these findings support a model in which combinatorial phosphorylation encodes specific tau proteoforms with functions that are tightly integrated with MT dynamics and cellular resilience.

## DISCUSSION

PTMs like phosphorylation diversify the functions of proteins. Tau is a striking example of PTM complexity, with over 80 potential phosphorylation and 40+ potential acetylation sites, as well as additional modifications including methylation, ubiquitination, succinylation, *O*-GlcNAcylation and others^14,51-53^; despite their abundance, these modifications are often studied in isolation. Our study examines the combinatorial landscape of tau phosphorylation in a physiological context to elucidate how its role is potentially disrupted in a pathological one.

While high levels of p-tau are most commonly described in the context of aggregates in disease, similar p-tau epitopes have been reported in a number of animal studies and recapitulated in our work here using PS19 mouse brains (**Fig. 1**). However, in the aged PS19 mouse, we found only a weak correlation between the relative abundance of a specific p-tau epitope and its presence in insoluble aggregates. For example, pS202/pT205 showed a large decrease in abundance in the aged brain compared to neonates but was strongly associated with insoluble tau; conversely, pT212/pS214 showed a large increase in abundance but was also highly associated with insoluble tau. Thus, while p-tau is generally enriched in insoluble fractions from aged PS19 mice, this raises the intriguing possibility that combinatorial phospho-regulation rather than individual phospho-epitopes may contribute to aggregation propensity in disease, as well as the extent to which phosphorylation at some epitopes largely occurs after aggregate formation.

To provide more insight, we turned to iNeurons, which we demonstrate are a powerful system for the study of endogenous tau phosphorylation both in the context of function and in disease. By top-down mass spectrometry of iNeuron-derived tau, we showed that endogenous fetal tau is extensively phosphorylated, with up to 21 phosphate groups per molecule – far exceeding prior estimates from fetal or AD brains^12,24,54,55^. Interestingly, a previous study similarly identified two pools of p-tau centered around ∼8 and ∼14 phosphates per tau molecule, by utilizing 2N4R overexpression in insect cells^24^. However, in that study and in contrast to our current work, the cells were pre-treated with phosphatase inhibitors prior to lysis to enrich p-tau; meanwhile, our results support a natural higher level of p-tau in iPSC-derived human neurons.

The use of iNeurons allowed us to perturb MT stability chemically in order to measure dynamic alterations to p-tau during MT destabilization and recovery. Importantly, we identified that specific combinations of phospho-sites are co-regulated, shifting in response to changes in MT stability and growth cycles (**Fig. 7**). When MTs destabilize, such as during the frequent collapse and regrowth cycles of axons, the first pool of p-tau species, phosphorylated at S202/T205, T212 and T217, are dephosphorylated presumably to enhance tau’s binding capacity to tubulin/MTs. Then, a second pool of p-tau species (i.e., pT231, pS235, pS262) is dephosphorylated to promote further MT stabilization. Based on our MT fractionation measurements for each phospho-epitope, the rapid dephosphorylation of the first set of sites (S202/T205, T212 and T217) may indicate a cellular response to increase the affinity of p-tau species which are already bound to MTs. Meanwhile, the slower dephosphorylation of the second set of phospho-epitopes (T231, S235 and S262) may represent dephosphorylation of a cytoplasmic fraction of tau, thus contributing to the overall population of dephosphorylated tau that is available to promote MT growth and repair. Among the phospho-sites measured in this study, dephosphorylation at sites T231, S235 and S262 appear to be the most critical for enhancing this interaction. Once MT stability is re-established, these same sites are re-phosphorylated: first, epitopes S202/T205, T212 and T217 are phosphorylated, possibly to facilitate dissociation of tau from MTs: then, tau is re-phosphorylated at sites that strongly inhibit tau’s binding to MTs (e.g., T231, S235 and S262) in order to maintain cytosolic pools of p-tau that are dissociated from MTs.

**Figure 7:**
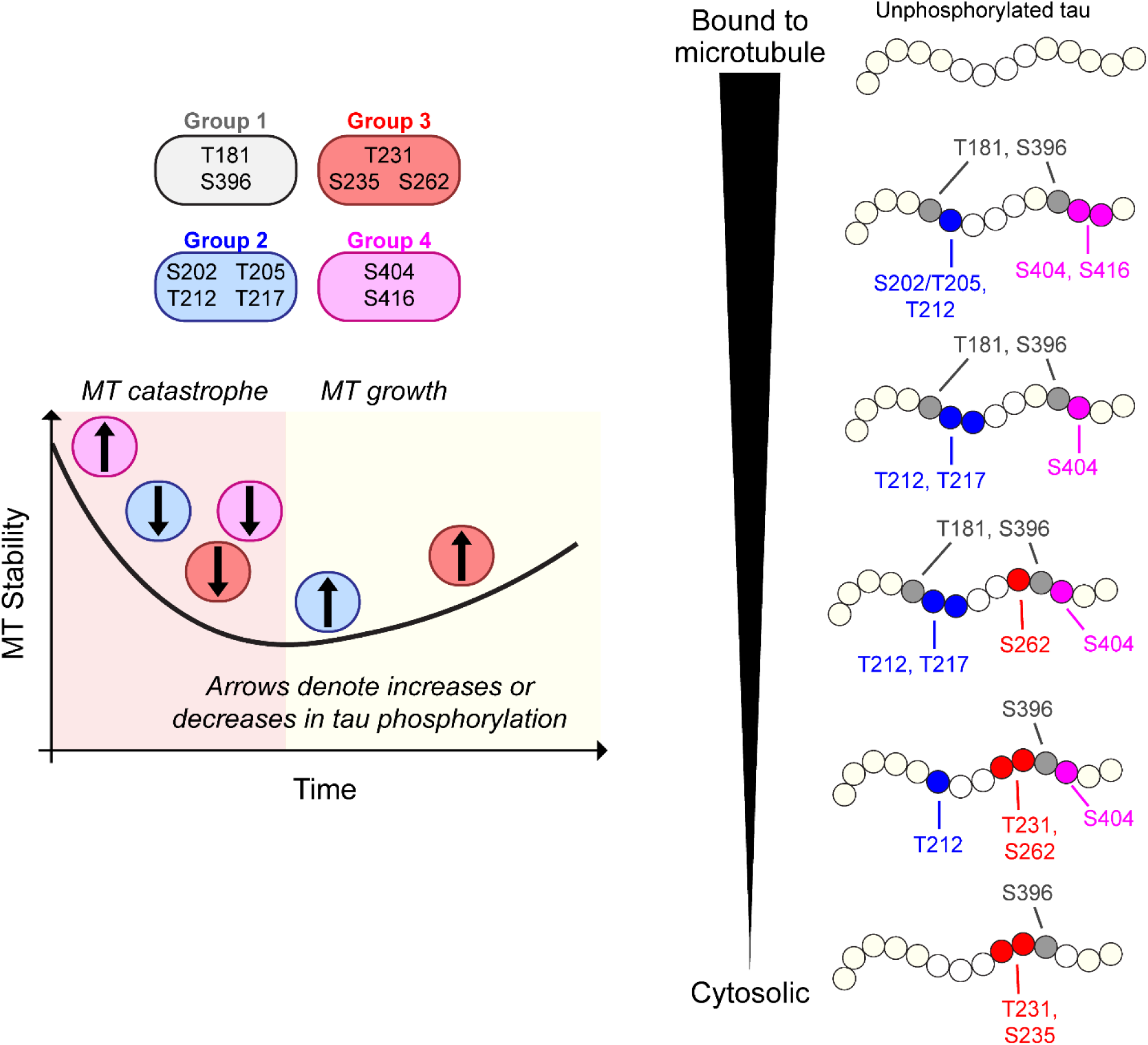
Schematic summary. Temporal changes to site-specific phosphorylated occur in specific combinations in response to MT collapse and growth. Based on co-occurrence linkage of phospho-epitopes, fractionation of MT-bound and cytosolic proteins, and changes to site-specific phosphorylated during MT collapse and recovery, we present predictions of the likelihood that certain combinatorial phospho-modifications to tau are found bound to or dissociated from MTs.

This paradigm contrasts sharply with the prevailing view in neurodegeneration in which increased tau phosphorylation is thought to drive MT destabilization. In fact, when we seeded iNeurons with human brain-derived tau aggregates, we observed an abundant presence of tau aggregates and globally elevated p-tau levels (**Fig. 2**), and thus a presumed loss of functional tau, but we did not detect a decrease in MT stability. Our findings question the “tau cascade” of MT destabilization by suggesting a more nuanced relationship: in physiological settings, tau phosphorylation responds to cytoskeletal changes and allows it to regulate MT stability, while in disease, tau phosphorylation broadly still responds to cytoskeletal changes, but tau becomes sequestered in aggregates and therefore is not available for function. In other words, tau may not simply stabilize MTs, but rather may maintain dynamic MT instability, consistent with recent findings that tau enhances tubulin incorporation preferentially at defect sites in the MT lattice^56^.

Our data suggest, but do not prove, that depletion of soluble tau through aggregation impairs tau’s ability to counteract MT destabilization. In seeded neurons, we observed increased phosphorylation across most epitopes tested, which was sustained at only a few sites during MT collapse, notably at epitopes S235 and S262. These two epitopes are among the most disruptive to tau-MT interactions, are highly sensitive to PP2a activity, and thus appear essential for regulating tau’s ability to stabilize MTs. Consistent with this, PP2a inhibition in unseeded neurons markedly blunted dephosphorylation at S235 and S262 during MT collapse, and resulted in MT destabilization comparable to levels seen in AD-seeded neurons. Given that S235 and S262 lie within or adjacent to the proposed fibril core of tau, reported by cryo-electron microscopy to span approximately residues 254-381 or more concisely from 306-378 (depending on the structural model)^57-60^, its apparent inaccessibility within aggregates may prevent normal PP2a-mediated dephosphorylation, locking tau in an elevated phosphorylated state that is unable to respond to MT stress.

Our findings point to site-specific, combinatorial layers of phospho-regulation that govern tau’s function. Our prior work with recombinant tau and phospho-mimics led us to propose a model whereby the contributions of individual phospho-modifications to disrupting interactions with MTs were relatively minor, but the accumulation of multiple modifications could significantly disrupt function^16^. Our work here refines that model to suggest that some individual phospho-sites are more disruptive than others are to MT-binding, but underscores the importance of multiple modifications in fine-tuning tau-MT interactions in a biological setting. Many questions remain about how combinatorial tau phosphorylation is regulated and how it influences tau’s diverse functions. Most of the pairwise and triplet phospho-combinations we identified are novel, leaving open how, when and where these phospho-proteoforms arise in neurons, and what roles they play beyond regulating tubulin/MT interactions. For example, we linked phosphorylation at S262 to phosphorylation at epitopes T212, T231, S235 and S396, all of which are highly abundant in AD brains^14^. Thus, the diversity of phospho-proteoforms may allow tau to carry out distinct cellular roles, including MT regulation, tubulin-actin crosslinking, vesicle trafficking, or synaptic modulation, depending on its modification profile.

Importantly, our study has limitations. Phospho-specific antibodies may vary in affinity and can be affected by neighboring modifications, which may lead to underestimation of phospho-occupancy at certain sites. For example, T181 phosphorylation frequently co-occurred with other phospho-sites while also exhibiting high levels of independent phosphorylation, yet we likely underestimated its occupancy at only ∼35%. Another limitation is that the use of kinase/phosphatase inhibitors and cytoskeleton-targeting small molecules are non-specific, and may affect tau indirectly, outside of physiological pathways that normally modify MT stability. Future work using genetic approaches to modify trans-regulators of tau-MT interactions, high-resolution top-down proteomics or genetic code-expansion strategies to incorporate site-specific PTMs, could more precisely resolve how phospho-proteoforms are regulated and function.

In summary, our work reframes several central assumptions in the tau field. We demonstrate that while single phospho-epitopes may be useful as biomarkers in tissue or CSF^5-8^, they are poor predictors of tau insolubility. Moreover, MT integrity can be maintained even in the presence of increased tau phosphorylation and accumulation of aggregates. These insights were made possible by leveraging iNeurons as a model of endogenous tau phosphorylation, allowing us to dissect combinatorial phospho-patterns in both healthy and disease-relevant contexts. We propose a mechanistic link between combinatorial tau phosphorylation and an impaired ability to counteract MT destabilization in the presence of tau aggregates, and that this phenotype is strongly correlated with elevated phosphorylation at epitopes S235 and S262. Our findings reveal that combinatorial tau phosphorylation patterns are tightly linked to MT dynamics and are organized in co-regulated clusters that may foreshadow aggregation behavior in disease. These insights reframe our understanding of tau regulation, emphasize the disruption caused by increased phosphorylation at S235 and S262 within tau aggregates, and highlight new avenues for exploring how phosphorylation contributes to tau function and dysfunction. Future studies may benefit from identifying specific phospho-proteoforms in physiological and pathological contexts that could reveal tau molecules that are unique to the disease state.

## METHODS

### Mouse lines and husbandry

Ethical Statement: All animal procedures including husbandry and experimental procedures, were performed according to NIH guidelines and approved by the Institutional Animal Care and Use Committee at Weill Cornell Medicine (Animal Welfare Federal Wide Assurance No: A3290-01; protocol number 2014-0015).

Mouse strains: PS19 mice^61^ overexpressing human tau (1N4R) carrying the disease-causing P301S mutation, driven by the mouse prion protein promoter (stock# 008169, Jackson Lab, ME) and wildtype (C57BL/6 J; Stock # 000664 Jackson Labs, ME) were kept in 12 h light/dark cycle in a temperature-controlled room with free access to water and food. PS19 transgenic mice (generated in C57BL/6 background, and inbred for >20 generations) were bred to wildtype mice of the same line to generate 50% transgenic and 50% wildtype offspring used in this study. Littermates were raised together and tissue were harvested in parallel.

### Mouse tissue harvesting

Mouse brains were dissected, washed once with ice-cold PBS, and cut into left and right hemispheres. The brains were flash-frozen in under 60 seconds post-decapitation. For post-natal brain dissections, one hemisphere was immediately flash-frozen in liquid nitrogen and stored at -80C for solubility assays. The other hemisphere was immediately resuspended to 10% (w/v) in cold PBS supplemented with protease and phosphatase inhibitors, mechanically homogenized using a dounce tissue grinder, and supplemented with 4x Laemmli buffer (supplemented with 10 mM DTT). The same homogenization protocol was used for embryonic brain dissections (E12). These samples were stored at -80C to be used for SDS-PAGE followed by immunoblot.

### Primary neuron cultures

Primary neurons from embryonic day 12 wildtype mice were cultured essentially as previously described^62^. Briefly, cortices were dissected in ice-cold HBSS followed by trypsinization (0.05% trypsin-EDTA) for 10 min at 37 °C. The brains were then triturated with a siliconized pipette and plated onto poly-L-lysine coated glass coverslips. Plating medium (DMEM, Thermo Fisher) containing 5 g/L glucose, 0.2 g/L NaHCO_3_, 0.1 g/L transferrin, 0.25 g/L insulin, 0.3 g/L L-glutamine, and 10% fetal bovine serum was replaced with growth medium (DMEM, Thermo Fisher) containing 5 g/L glucose, 0.2 g/L NaHCO_3_, 0.1 g/L transferrin, 0.3 g/L L-glutamine, 5% fetal bovine serum, 2% B-27 or N21-MAX supplement, and 2 μM cytosine arabinoside 2 days after plating. On day *in vitro* 8, cultured neurons were washed twice with cold PBS and fixed with 4% PFA (in PBS) for 20 minutes at room temperature, followed by immunofluorescence.

### Generation of tau^KD^ human iPSCs

Human WTC11 iPSCs stably-expressing Ngn2 (under a dox-inducible promoter) and dCas9-KRAB_(KOX1)_ were transduced with sgRNAs targeting *MAPT* followed by puromycin selection. Three sgRNAs were tested independently: 1) 5’GTGTTGGTGCCGGAGCTGGT, 2) 5’GGGCTGCAGTCGAGAGTGAA, and 3) 5’GGAAGCCCCAGTCTGCGGAG. Non-targeting control (NTC) sgRNA: 5’ GCCGAGCGGCAGGGTGCCGC. The first sgRNA resulted in stable knockdown of tau protein for at least 45 days post-differentiation while the other two sgRNAs resulted in only partial tau knockdown. The described experiments were performed using iPSCs generated from the first sgRNA, referred to as tau^KD^.

### Lentivirus production and transduction

Lentivirus was generated as previously described^62^. Briefly, HEK293T cells were co-transfected at low passage with 1.06μg of pMDLG (Addgene plasmid# 12251), 0.57μg of pMD2.G (Addgene plasmid# 12259), 0.4μg of pRSV-Rev (Addgene plasmid# 12253) plasmid, and 1.6μg of the lentiviral plasmid, using 1:3 ratio of total DNA to PEI per well of a 6-well plate. The culture medium was replaced after 6 hours with mTesR+ (for iPSC transductions) or neurobasal media (for neuronal transductions). The supernatant was collected after 48h and passed through a 0.45 μM polyether sulfone filter to remove any cell debris. Large batches of virus were made together and small aliquots were stored at -80°C, to be thawed only once.

iPSCs were transduced with lentivirus containing guides against *MAPT* or NTC followed by puromycin selection (1 µg/mL) 48 hours-post transduction for 72 hours (until control, non-transduced cells were 100% dead) to generate tau^KD^ iPSCs.

iPSC-derived tau^KD^ neural progenitors were transduced with lentivirus containing tau constructs at 72 hours post-differentiation using the doxycycline-induced Ngn2 cassette. Neural progenitors were lifted with StemPro Accutase Cell Dissociation Reagent (5 min, 37C), pelleted and resuspended in neuronal differentiation media. Neural progenitors were plated at 0.5×10^6^ cells in a single well of a 6-well plate. Lentiviruses containing either FUW-0N3R^myc^ or FUW-0N3R^EGFP^ were added to the media upon re-plating. We did not observe any silencing of exogenously expressed tau in differentiated neurons using the UbC promoter.

### iPSC culture and neuronal differentiation

Human iPSC cultures and neuronal differentiation protocols were performed as previously described with minor modifications^63^. CRISPRi-iPSCs expressing inducible Ngn2 from AAVS1 safe harbor locus were maintained in mTesR1+ media with mTesR1+ supplements. iPSCs were plated on hESC Matrigel and maintained in mTesR1+ media (Stem Cell Technologies). iPSCs were differentiated to glutamatergic neurons following established protocols^63^. iPSCs were lifted and singularized using StemPro Accutase (5 min, 37C) followed by neutralization with mTesR1+ media. Cells were gently pelleted for 5 minutes at 300*g*, then resuspended to 0.7×10^6^ cells/mL in neuronal induction media: KnockOut DMEM/F-12 (Gibco# 12660012) + 1x NEAA (Gibco#11140050) + 1x GlutaMAX (Gibco# 35050-061) + 10 nM Y27632 ROCK inhibitor (Tocris# 125410) + 1x N2 max (Gibco #17502-048) + 2 µg/mL doxycycline hydrochloride (Sigma# D9891). A complete media change was made the following day using the same media. The next day, another complete media change was made but without ROCK inhibitor. Neural progenitors were lifted and dissociated with StemPro Accutase 72-hours post neuronal induction, pelleted (5 min, 300*g*), and re-plated at 0.5×10^6^ cells per well of a 6-well plate that was first coated overnight with poly-L-ornithine hydrobromide (0.1 mg/mL in borate buffer pH 8.5; Sigma, #P3655). The replating maturation media contained: Neurobasal-A (Gibco# 10888-022) + 1x NEAA + 1xGlutaMAX + 10 nM Y27632 ROCK inhibitor + 2 µg/mL doxycycline hydrochloride + 1x B-27 plus (Gibco# A3582801) + CultureOne Supplement (Gibco #A3320201) + 10 ng/mL BDNF (PeproTech #450-02) + 10 ng/mL NT-3 (PeproTech #450-03) + 1 µg/mL laminin (Gibco# 23017-015) + 200 µM L-ascorbic acid (Sigma #A8960). 50% media changes were made every 7 days with the same media lacking ROCK inhibitor and doxycycline. iNeurons were harvested at the time points indicated within each figure and figure legend.

WTC11 iPSC-derived neurons were used for all the experiments except for in tau seeding experiments (‘Figure 2’), which utilized KOLF2.1J iPSC-derived neurons. Differentiation protocols were essentially identical except the neural progenitor phase was extended by an additional 4^th^ day which was also supplemented with the anti-mitotic 5-fluorodeoxyuridine (5-FdU).

### Top-down mass spectrometry

Top-down mass spectrometry (TD-MS) of tau proteoforms was performed essentially as previously described^64^. The harvested iNeurons (8 million cells, DPI 30) were suspended in 500 μL of homogenization buffer composed of Ham’s F12 medium supplemented with 1× Halt Protease and Phosphatase Inhibitor, 5 mM EDTA, and 0.45% CHAPS, and left in 4 °C for half an hour for cell lysis. Afterwards, the cell lysate was subjected to sonication on ice for 20 seconds at 40% power, using a pulse cycle of 2 seconds on and 3 seconds off. The lysate was then centrifuged at 17,000*g* for 30 minutes at 4°C to separate cell debris, and supernatants were collected for tau IP. Tau antibody-conjugated beads were prepared by incubating 10 μg each of HT-7 and Tau-5 antibodies with 50 μL of Protein A/G beads at room temperature for 10 minutes. The prepared conjugated beads were added to 500 μL of the cell lysate and incubated overnight at 4°C with end-to-end rotation. Afterwards, the beads were subjected to a series of wash steps: twice with 500 μL homogenization buffer, twice with a 1:1 mixture of homogenization buffer and water, and three times with 500 μL of water. Finally, the tau proteins were eluted from the beads incubation in 150 μL of 10% formic acid at room temperature for 30 minutes. The eluates were flash-frozen and stored at -80 °C for further analysis.

Individual ion mass spectrometry (I^2^MS) is an advanced top-down MS technique that enables direct mass readout of complex tau proteoforms at the intact level^65^. The SampleStream platform (Integrated Protein Technologies) was interfaced with a Q Exactive HF mass spectrometer (Thermo Fisher Scientific) using the Ion Max source equipped with a HESI II probe to provide online buffer exchange and I^2^MS analysis. Triplicate I^2^MS runs were performed for the iNeuron-derived tau IP sample. For each run, 50 μL of immunoprecipitated tau samples were cleaned and buffer-exchanged into a solution of 0.2% (v/v) formic acid and 30% (v/v) acetonitrile in water, followed by direct injection into the mass spectrometer at a flow rate of 1.6 μL/min for I^2^MS analysis. On the mass spectrometer, automated Ion Control (AIC) was utilized to dynamically adjust the injection time based on ion signal density, ensuring optimal detection of individual ion signals. Additional mass spectrometry settings included an Orbitrap central electrode voltage of 1 kV, trapping gas pressure of 0.3, m/z range of 650-2,500, a resolution of 120,000 at m/z 200 (corresponding to a 1 s transient time), a spray voltage of 3.1 kV, in-source CID energy of 15 eV, and a source temperature of 320 °C. I^2^MS data were acquired over a 50-minute period per run, totaling approximately 3,000 scans.

All the I^2^MS transient files were converted to .storx files and processed using STORIboard (https://www.proteinaceous.net/storiboard) with the voting charge assignment algorithm. After processing, .mzML files were generated, presenting the mass domain spectrum of each run. The mass spectra were annotated by TDValidator (Proteinaceous, https://www.proteinaceous.net/tdvalidator). Tau 0N3R proteoforms were matched to spectral peaks based on intact mass, using a mass accuracy window of ±50 parts per million (ppm) and a mass tolerance of 1.8 Da. The 0N3R proteoforms were quantified by their relative abundance.

### SDS-PAGE and quantitative immunoblotting

Proteins were separated by SDS-PAGE for immunoblotting as previously described^62^. Briefly, 8-15% Laemmli gels (10.3%T and 3.3%C) were used to separate proteins on Bio-Rad apparati. 2.5 mM final DTT was used in SDS Laemmli buffer unless indicated otherwise. Proteins were transferred onto nitrocellulose (pore-size = 0.45 μm; GE Healthcare) and blocked with Every Blot Blocking Buffer (Bio-Rad). Immunoblotting was performed by incubating the blocked membranes with primary antibodies in blocking buffer for overnight incubation. Following 5 washes with TBS-Tween 20 (0.1%), blots were incubated with secondary antibodies (goat anti-rabbit conjugated to IRDye 680RD or 800CW; LI-COR) at 1/10,000 in blocking buffer for 1-2 hours. Immunoblots were washed 5x with TBS-T and dried, then scanned on an Odyssey CLx (LI-COR) and quantified using Image-Studio software (LI-COR).

### Triton-X solubility assay of PS19 transgenic mouse brains

Mouse brains from PS19 mice were dissected on ice at either post-natal day 0 or at 9-months. Brain lysates were subject to Triton-X (Tx) solubility as previously described^66^. Briefly, brains were rinsed twice with ice cold PBS, then homogenized in 0.1% Tx-100 in PBS supplemented with protease and phosphatase inhibitors. Samples were rotated for 30 min at 4C and then sonicated 10x at 10% power (0.5 sec on, 1 sec off). The tissue was then centrifuged at 17,000*g* for 90 min at 4C. The supernatant (soluble fraction) was collected, and the pellet (insoluble fraction) was resuspended in an equal volume of lysis buffer as the supernatant. Each sample volume was doubled using 4x Laemmli buffer (supplemented with 10 mM DTT) and sonicated again: 20x at 10% power (0.5 sec on, 1 sec off). The samples were then boiled at 95C for 20 min before loading equal volumes of each fraction on SDS-PAGE, followed by immunoblot for tau. Approximately two-fold more tissue levels were loaded for 9-month-old samples compared to neonates (as indicated by β-actin immunoblot) to improve detection of lowly-detected phosphorylation epitopes, particularly at sites pS202/pT205, pT231 and pS262. Insoluble tau was calculated as a percentage of total protein per sample and/or normalized to β-actin, as indicated.

### *0N3R* recombinant aggregation and quantification

Aggregation of 0N3R was prepared as previously described^67^. Briefly, 50 µM tau was supplemented with 500 µM DTT in a sample volume of 100 μL of 1x PBS buffer, pH 7.4. The sample was then left on an IKA MS 3 control shaker with continuous orbital shaking at 1000 rpm at 37C. Following a six-day aggregation period, the aggregates were pelleted by centrifugation at 17,700*g* for 90 minutes. The supernatant was then separated from the pellet, and the pellet was resuspended to 100 µL. The supernatant and pellet samples were boiled for 20 minutes and chilled on ice before an SDS-page gel was conducted to calculate the approximate amount of protein in the pellet.

### TEM Imaging

The same tau 0N3R pellet was imaged using transmission electron microscopy (TEM) (Talos L120C TEM, ThermoFisher). The fibrils (5 μL of 10 μM) were added to freshly glow-discharged carbon coated film 300 mesh grids (Electron Microscope Sciences) and incubated for 1 minute. A wash with 5 μL water for 30s was performed to remove excess buffer salts. To negative stain, the sample was incubated with 5 μL uranyl acetate (2% in water) for 1 minute. Between each step, excess buffer was absorbed with filter paper. The grids were dried under vacuum pump for 2 minutes before imaging. The 0N3R pellet sample was imaged with an accelerating voltage of 120 kV and a magnification of 22000x.

### AD case selection and tau seeds extraction

The AD patient was pathologically diagnosed as previously described^68^. The case was pathologically confirmed as late-stage AD without comorbidities of Lewy body and TDP-43 pathologies based on immunohistochemical staining (data not shown). Tau seeds were sequentially extracted from the frontal cortex using a previously described protocol^69^. Grey matter and white matter were separated using a surgical blade, and only grey matter was used in the extractions. Brain homogenate was prepared in nine volumes (v/w, mL/g) of PHF buffer (10 mM Tris, 10% sucrose, 0.8 M NaCl, 1 mM EDTA, pH 7.4) with 0.1% sarkosyl, proteinase inhibitor and phosphatase inhibitor in a glass dounce homogenizer and spun at 10,000*g* for 10 min at 4C. Supernatant (supernatant 1) was collected, sarkosyl was added to 1% final concentration, incubated in a beaker with stirring for 1.5hr at RT, and followed by 150,000*g* spin for 75 min at 4C. The pellet was collected, briefly washed with PBS to remove myelin, and resuspended in PBS (as pellet 1). To remove the sarkosyl, pellet 1 was spun at 150,000*g* for 75 min, and the resulting pellet was collected as pellet 2. Pellet 2 was resuspended in PBS, thoroughly sonicated, and spun at 10,000*g* for 10 min at 4C. The supernatant was collected as the enriched tau seeds.

### Tau seeding and aggregate detection

Protocols for AD-derived tau seeds were adapted from previously established protocols^70^. KOLF2.1J-NGN2 iNeurons were plated at either 0.2×10^6^ or 0.05×10^6^ cells per well for biochemistry or immunocytochemistry experiments, respectively. Three weeks post-NGN2 induction, AD-derived seeds (0.1 µg/mL final) or recombinant 3R seeds (1.28 µg/mL final) were diluted in 100 µL total of neuron maturation media. Seeds were sonicated on high in a temperature-regulated water bath for 20 minutes at 10C for pulses of 30 seconds on, 30 seconds off. The sonicated seeds were immediately transferred to ice and further diluted to a total 720 µL in neuron maturation media. 20 µL of seed solution was added dropwise in a circular motion to each treated well, triturated gently 5x, and then the plate was gently rocked back and forth. The seed solution was triturated 5x between each treated well with a fresh tip. iNeurons were then maintained as described in the ‘*iPSC culture and neuronal differentiation*’ section using a 50% change of neuron maturation media with supplements (as indicated) every 7 days. For biochemistry experiments, iNeurons were treated at 4-weeks post-seeding with either DMSO or 10 µM nocodazole for 90 minutes, as described in the ‘*Cytoskeleton manipulation by pharmacological reagents’* section.

### Immunofluorescence in primary mouse neurons and iNeurons

Immunofluorescence was performed essentially as previously described^62^. Cells were plated and maintained on tissue-culture treated and coated glass coverslips (poly-L-laminin for primary neurons, poly-L-ornithine for iNeurons). Cells were washed twice with PBS + 1 mM MgCl_2_ and fixed in 4% paraformaldehyde (PFA) for 20 min at room temperature. The PFA was washed out twice with PBS + 1 mM MgCl_2_, then the cells were permeabilized for 5 minutes with 0.1% Tx-100 in PBS-T, followed by incubation with blocking buffer (PBS-T + 5% BSA) for 1 hour at room temperature. Cells were incubated overnight at 4C in primary antibody: β-tubulin III rabbit polyclonal (1:250), β-tubulin III mouse monoclonal (1:250), α-tubulin 1a mouse monoclonal (1:250), detyrosinated α-tubulin rabbit polyclonal (1:250).

For seeding experiments, iNeurons were fixed in ice-cold methanol at -20C for 15 minutes to extract soluble proteins, followed by 1 hour incubation with blocking buffer (3% BSA and 3% goat serum), followed by overnight anti-AT8 (1:1000) staining at 4C for detection of p-tau aggregates, or anti-β-tubulin III mouse monoclonal (1:250) for detection of polymerized tubulin.

### Pharmacological reagents

The following pharmacological reagents were utilized throughout this study: Nocodazole (10 µM final; 12-281-0), demecolcine (1 µM final; D7385; lot #102753904), paclitaxel (10 µM final; T1912; lot #1003615845), latrunculin A (2 µM final; L5163; lot #0000373456), LB-100 (20 µM final; 29105; lot #0613520-17), Calyculin A (20 nM final; C5552; lot #0000366393), AKT1 inhibitor VIII (20 µM final; 14870; lot #0613313-34), Erk1/2 inhibitor I (10 nM final; A1176841; lot #HA2), GSK3β BIO-1 inhibitor (5 µM final; 13123; lot #0586619-49).

### Cytoskeleton manipulation by pharmacological reagents

iNeurons (*DPI 21*) were treated with either DMSO, nocodazole (10 µM), demecolcine (1 µM), paclitaxel (10 µM) or latrunculin A (2 µM) for 90 minutes in complete neuronal media. Cells were washed twice with PBS at room temperature and harvested for Western blot in ice-cold RIPA buffer supplemented with protease and phosphatase inhibitors.

### Pharmacological microtubule catastrophe by nocodazole treatment

Nocodazole was resuspended in DMSO to 10 mM, aliquoted and stored at -20C. DPI 25 iNeurons (0.5×10^6^ per well of a 6-well plate for immunoblot, 0.4×10^5^ per well of a 24-well plate for immunofluorescence) were washed once with ice-cold PBS for 5 min to facilitate MT catastrophe, and then treated with 10 µM nocodazole (final) in pre-warmed complete neuronal media for either 15 minutes or 90 minutes at 37C. For MT recovery experiments, nocodazole was washed out by aspirating out the media and performing three washes with pre-warmed neuronal media without nocodazole. After the third wash was removed, the cells were incubated at 37C in complete neuronal media without nocodazole for either 1 hour, 3 hours or 6 hours to allow for MT recovery. At the final time point, for immunoblot, the cells were washed twice with room temperature PBS then lysed and scraped in ice-cold RIPA buffer supplemented with protease inhibitors and phosphatase inhibitors on ice. Following 10 min of rotation at 4C, debris were pelleted at 17,000*g* for 10 min at 4C, the supernatant was collected and supplemented with 4x Laemmli buffer + 10 mM DTT, boiled for 20 min at 95C, then subject to SDS-PAGE followed by immunoblot. For immunofluorescence, the cells were washed twice with 0.1M PHEM buffer (pH 7.2) at room temperature and fixed for 15 min in 4%PFA/0.25%GA in 0.1M PHEM buffer (pH 7.2) at room temperature. The fixative was washed out 3x with PHEM buffer, and immunofluorescence was performed as described above except PBS was replaced with PHEM buffer for all steps including washing and blocking buffer (5% BSA in PHEM-T). Stable microtubules were detected by anti-acetylated α-tubulin and anti-detyrosinated α-tubulin antibodies.

### Kinase/phosphatase inhibition

iNeurons (*DPI 30*) were treated for 90 minutes with nocodazole (10 µM) and either LB-100 (20 µM) or Calyculin A (20 nM). Cells were either washed and collected for Western blot at this point, or the nocodazole and phosphatase inhibitors were washed out 3x with pre-warmed complete media (lacking nocodazole and phosphatase inhibitors) and allowed to recover at 37C for 3 hours prior to harvesting. For kinase inhibition, neurons were treated for 90 minutes with 10 µM nocodazole followed by 3x washout, followed by incubation with AKT1 inhibitor VIII (20 µM) for 3 hours at 37C. At this final timepoint, cells were then harvested, as described above, for Western blot.

### Intact microtubule fractionation from iNeurons

Intact microtubules were fractionated from soluble cytosolic proteins using tau^KD^ iNeurons expressing FUW-0N3R^myc^ at 12 days post-neuronal induction following established protocols^71^. Neurons were plated at 0.5×10^6^ cells in a single well of a 12-well PLO-coated plate.

Fresh microtubule stabilization buffer (MSB) was made containing 100 mM PIPES (Millipore Sigma #P1851) + 1 mM MgSO_4_ (M2643) + 2 mM EGTA (E3889) + 0.1 mM EDTA (E8008) + 2 M glycerol (G9012). The pH was adjusted to 6.75 with KOH (221473) to fully dissolve the PIPES. 1 mM PMSF (10837091001) + 1x protease inhibitor (P8340) + 1x Halt phosphatase inhibitor cocktail (78420) were added fresh immediately before the assay. MSB was pre-warmed to 37C.

The media was aspirated, and the cells were gently washed once with MSB. Next, the cells were incubated with 300 µL of pre-warmed MSB containing 1% Tx-100 for 5 min at 37C. The plate was gently tilted after the addition of the buffer. After 5 minutes, the buffer was collected (and placed on ice) and the cells were washed with 300 µL of the same buffer and combined with the initial collection, termed the cytosolic fraction. Cells with intact cytoskeletons remained adhered to the well. These were scraped in 600 µL MSB containing 1% Tx-100 and placed on ice (intact MT fraction). Following the extractions, the two fractions were brought to equivalent volumes in MSB + 1% Tx-100. 200 µL 4X Laemmli SDS loading buffer (supplemented with 10 mM DTT) was added to each sample. Samples were boiled for 20 min at 95C and immediately subject to separation by SDS-PAGE for immunoblot.

The same protocol was used for immunofluorescence except cells were either incubated for 5 min at 37C in MSB (no cytosolic extraction) or MSB + 1% Tx-100 (cytosolic extraction), followed by a single wash in the same buffer and then fixation by 4% PFA in PBS for 20 min at room temperature.

Anti-detyrosinated α-tubulin was used as a marker for intact MTs, and anti-NSE was used as a cytosolic/membrane marker.

### Purification of recombinant tau wildtype and phospho-mimic proteins

Recombinant tau proteins were expressed in BL21(DE3) cells and purified as previously described with minor modifications^16^. *MAPT* was cloned into the pET-HT vector with a N-terminal 6xHis-TEV fusion, and transformed into BL21(DE3) competent cells. Overnight starter cultures were incubated in 500 mL LB broth at 37C until OD600=0.6, at which time protein expression was induced with 1 mM isopropyl-β-D-1-thiogalactopyranoside (IPTG) at 18C for 16 hours. Bacteria were then pelleted (4000*g* for 20 min at 4C) and resuspended in cold lysis buffer (20 mM Tris pH 8.0, 150 mM NaCl, 1 Roche EDTA-free protease inhibitor tablet, 1 mg/mL lysozyme, 20 mM imidazole, 1 mM PMSF, 1x DNase, 1 mM MgCl_2_). Cells were rotated at 4C for 30 minutes followed by sonication on ice (2 minutes; 1 sec on, 2 sec off) and centrifuged again (20,000*g*, 30 minutes at 4C) to remove cellular debris. The supernatant was subject to affinity capture using Ni-NTA resin (Fisher; PI88222) for 1 hour, rotating, at 4C. Unbound proteins were washed away with 6 column volumes of buffer containing 20 mM Tris pH 8.0 + 150 mM NaCl + 20 mM imidazole, then eluted in the same buffer containing 300 mM imidazole. The eluted 6xHis-TEV-tau protein was buffer exchanged to the previous buffer (20 mM Tris pH 8.0 + 150 mM NaCl + 20 mM imidazole) using a 10 kDa MWCO Amicon Ultra-15 Centrifugal Filter Unit (Millipore #UFC9010), concentrated to ∼1.5 mL, and incubated with super TEV protease-6xHis (1:300 enzyme to tau molar ratio) and 1 mM DTT, rotating at 4C overnight. The following day, the mixture was diluted 15-fold in 20 mM Tris pH 8.0 + 150 mM NaCl + 20 mM imidazole to dilute the DTT and reverse-Ni NTA affinity capture (1 hour, rotating at 4C) was used to remove the TEV-6xHis protease from the mixture. The flowthrough (containing tau) was collected and buffer exchanged into 20 mM Tris pH 8.0 + 150 mM NaCl + 1 mM EDTA + 1 mM TCEP, concentrated to ∼2 mL and injected onto a HiLoad 16/600 Superdex 200 PG column for size exclusion in the same buffer. Fractions containing tau were identified by UV-absorption at 280 nm. Fractions containing >95% pure tau were pooled, concentrated to ∼100 µM (determined by NanoDrop), aliquoted, flash frozen in liquid N_2_ and stored at -80C to be used for a single freeze-thaw cycle.

Tau phospho-mimic mutation sequences are as follows: Thr181Glu (ACA>GAA): Thr217Glu (ACC>GAA): Thr231Glu (ACT>GAA): Ser235Glu (TCG>GAG): Ser262Glu (TCC>GAA): Ser396Glu (TCG>GAG): Ser404Glu (TCT>GAA).

### Purification of tubulin from bovine brains

Tubulin was purified from fresh bovine brains as previously described^72^ and flash frozen in BRB80 buffer (80 mM PIPES + 1 mM MgCl_2_ + 1 mM EGTA, pH 6.8). Frozen aliquots were thawed on ice and clarified by centrifugation at 100,000*g* for 6 min at 4C. The supernatant was collected, and the concentration of tubulin was estimated using NanoDrop. Clarified samples were used immediately for the MT polymerization assay, and fresh aliquots were clarified for each replicate.

### Microtubule polymerization assay by turbidity

Polymerization of tubulin was monitored by light scatter at 340 nm as previously described with minor adjustments^15^. Freshly clarified tubulin (20 µM final) was incubated with tau protein (10 µM final) in cold BRB80 buffer supplemented with fresh 1 mM DTT. A 250 µL reaction was made on ice and 100x fresh GTP (1 mM final) was added and mixed by pipette trituration on ice. After 5 minutes, 200 µL were quickly transferred to a pre-warmed 96-well black-walled clear bottom plate and the reaction was monitored at 37C for 45 minutes on a M1000 Tecan spectrophotometer. 9 flash measurements were made and averaged per well per timepoint throughout the kinetic measurement. Each replicate of 0N3R tau wildtype and phospho-mimics were measured in parallel. Three independent measurements were made and compared to their respective wildtype measurements for each sample, to account for read-to-read variability. Afterwards samples were returned to ice for 5 min to confirm that the samples could be depolymerized (thus eliminating concern for potential protein aggregation). Polymerization curves were corrected by subtracting a plate blank, a buffer blank and a tubulin + GTP (no tau) measurement from each sample. Curves were then normalized to the maximum data point for each sample. The time to 50% V_max_ was compared to wildtype for each isoform and averaged for final measurements.

### Immunoprecipitation of phospho-tau

Tau^KD^ iNeurons were re-plated at 2×10^7^ cells/plate on a PLO-coated 15-cm dish and transduced with FUW-0N3R^EGFP^. At 32 days post-induction, iNeurons were washed twice with room temperature PBS and then harvested in the dish in 8 mL of ice-cold RIPA buffer (supplemented with 2x protease inhibitors and 2x phosphatase inhibitors). The cells were transferred to a conical tube and rotated at 4C for 10 minutes, followed by pelleting of debris by centrifugation at 17,000*g* for 10 min at 4C. The supernatant was collected and divided into 8 equal tubes (1 mL each, ∼2.5×10^6^ cells per tube) for phospho-tau immunoprecipitation. 50 µL of lysate was collected from each tube and was supplemented with 50 µL of 4x Laemmli buffer (supplemented with 100 mM DTT, and boiled for 15 min at 95C before SDS-PAGE) for the input lanes. Phospho-tau antibodies were added to the remaining lysates and rotated at 4C overnight: pT181 mouse monoclonal (10µg), AT8 mouse monoclonal (10µg), pT212 rabbit polyclonal (10µg), pT217 rabbit polyclonal (10µg), pT231 mouse monoclonal (10µg), pS235 mouse monoclonal (15µg), pS396 rabbit polyclonal (5µg) or non-specific mouse or rabbit IgG (for IgG control). The following morning, either Protein A (Neta/Sigma, GE28-9513-78) or Protein G (Neta/Sigma, GE28-9513-79) magnetic sepharose beads were washed 3x with ice-cold RIPA buffer and incubated with each immunoprecipitation for 2 hours, rotating at 4C. The beads were then gently pelleted and washed 3x with ice-cold RIPA buffer using a DynaMag-2 Magnet rack (Fisher, 12-321-D). Affinity captured proteins were eluted into 170 µL of 2x Laemmli buffer (supplemented with 50 mM DTT) at 95C for 15 minutes. Samples were subject to SDS-PAGE followed by co-blotting for phospho-tau epitopes.

### Calculation of phospho-tau abundance and co-occurrence

Affinity-captured phospho-tau was assumed to be monomeric and soluble due to the presence of deoxycholate, triton-X and SDS in the lysis and wash buffers, and therefore co-blotted phospho-tau was assumed to be covalently-modified tau rather than the result of tau multimerization. Volumes per well for each measurement for SDS-PAGE: input (5µL, representative of ∼6.25×10^3^ cells, or 2.24% of the IP lane), IgG control or eluted phospho-tau (20µL, representative of ∼2.79×10^5^ cells). Signal intensities in the input lane were back-calculated to 100% of the IP lane (by multiplication by 44.7.

#### To calculate phospho-tau abundance

The efficiency of phospho-tau that was affinity-captured by each phospho-specific antibody was calculated using immunoblot:

A. Efficiency of phospho-tau capture = Intensity of p-tau IP lane / Back-calculated intensity of p-tau input lane.

Next, the percentage of total tau that was recovered within each affinity capture was calculated. This number represents the percentage of phospho-tau that was recovered from the total tau lysate.

B. Efficiency of total tau capture = Intensity of total tau IP lane / Back-calculated intensity of total tau input lane.

Lastly, phospho-tau abundance was estimated by using a ratio of ‘B / A’. In other words, of the A% of phospho-tau that was recovered, B% of it was positive for the given phospho-tau epitope. For example, for a given replicate, if 2.57% of pT181 was recovered (‘A’), and this was equal to 1.03% of total tau recovery (‘B’), then ‘B / A’ = 1.03% / 2.57% = 39.9% of total tau is phosphorylated at T181. Abundance was calculated using the average of a technical repeat for each replicate, and then averaged across all replicates: pT181 (n=4), AT8 (n=5), pT212 (n=4), pT217 (n=6), pT231 (n=6), pS235 (n=6), pS396 (n=4).

#### To calculate phospho-tau co-occurrence

The same logic as above was used to calculate phospho-tau co-occurrence:

C. Efficiency of phospho-tau co-occurrence = Intensity of p-tau co-blot in IP lane / Back-calculated intensity of co-blotted p-tau in input lane.

Co-occurrence was calculated using a ratio of ‘C / A’. In other words, of the A% of phospho-tau that was recovered, C% of it was positive for the given co-occurring phospho-tau epitope. For example, for a given replicate if 0.052% of AT8 was recovered (‘C’) from the pT181 immunoprecipitation (A = 2.57%), then ‘C / A’ = 0.052% / 2.57% = 20.1% of pT181 molecules carry a phospho-modification at the AT8 epitope.

Recovery of tau phosphorylated at Ser235 was too inefficient to reliably quantify co-occurrence of phosphorylation; therefore, these calculations were omitted.

Combinations of three co-occurring phospho-epitopes were predicted by minimum linkage analysis, which ranks trios based on their weakest link by accounting for co-occurrence in both directions. In other words, low P(A, B) will penalize the score even if P(A, C) + P (B, C) is high.

### Antibody list

| Antibody | Host Species | Manufacturer | Catalog # | Working Dilution |
| --- | --- | --- | --- | --- |
| Tau5 (pan-tau) | Mouse | Invitrogen | MA5-12808 | 1:1000 (WB) |
| Tau5 (pan-tau) | Rabbit | Novus | NB015844 | 1:1000 (WB) |
| HT7 (pan-tau) | Mouse | Invitrogen | MN1000 | For IP only |
| AT270 (Tau pT181) | Mouse | Invitrogen | MN1050 | 1:1000 (WB) |
| AT8 (Tau pS202/pT205) | Mouse | Invitrogen | MN1020 | 1:1000 (WB)<br>1:1000 (ICC) |
| Tau pT212 | Rabbit | Invitrogen | 44740G | 1:1000 (WB) |
| Tau pS214 | Rabbit | Invitrogen | 44742G | 1:500 (WB) |
| AT100 (Tau pT212/pS214) | Mouse | Invitrogen | MN1060 | 1:1000 (WB) |
| Tau pT217 | Rabbit | Invitrogen | 44-744 | 1:1000 (WB) |
| AT180 (Tau pT231) | Mouse | Invitrogen | MN1040 | 1:1000 (WB) |
| Tau pS235 | Mouse | Millipore Sigma | MABN2275 | 1:500 (WB) |
| Tau pS262 | Rabbit | Invitrogen | 44-750G | 1:500 (WB) |
| Tau pS324 | Rabbit | Abcam | ab109404 | 1:500 (WB) |
| Tau pS356 | Rabbit | Invitrogen | 44-751G | 1:500 (WB) |
| Tau pS396 | Rabbit | Invitrogen | 44-752G | 1:1000 (WB) |
| Tau pS404 | Rabbit | Invitrogen | 44-758G | 1:1000 (WB) |
| Tau pS416 | Rabbit | Invitrogen | PA1-4719 | 1:1000 (WB) |
| Tau pS422 | Rabbit | Invitrogen | 44-764G | 1:1000 (WB) |
| NeuN | Rabbit | Invitrogen | PA5-74899 | 1:1000 (WB) |
| GAPDH | Rabbit |  |  | 1:1000 (WB) |
| Tau 3R | Mouse | Millipore Sigma | 05803MI | 1:1000 (WB) |
| β-tubulin III | Mouse | R&D Systems | MAB1195 | 1:1000 (WB)<br>1:250 (ICC) |
| β-tubulin III | Rabbit | Invitrogen | PA5-86069 | 1:250 (ICC) |
| α-tubulin 1A | Rabbit | Abcam | ab18251 | 1:1000 (WB) |
| Detyrosinated α-tubulin | Rabbit | Millipore Sigma | AB3201 | 1:1000 (WB)<br>1:250 (ICC) |
| Acetylated α-tubulin (aK40) | Mouse | Invitrogen | 6-11B-1 | 1:1000 (WB) |
| β-actin | Mouse | Invitrogen | MA1-744 | 1:1000 (WB) |
| Hsp70 | Mouse | Invitrogen | MA3-007 | 1:250 (WB) |
| NSE | Rabbit | Abcam | ab53025 | 1:1000 (WB) |
| MAP1A | Rabbit | Invitrogen | PA558642 | 1:1000 (WB) |
| MAP1B | Rabbit | Invitrogen | PA578052 | 1:1000 (WB) |
| MAP2 | Rabbit | Millipore Sigma | AB5622 | 1:1000 (WB) |
| SV2a | Rabbit | Thermo Fisher | PA5-115990 | 1:1000 (WB) |
| pAkt1 (pS473) | Rabbit | Invitrogen | 44621G | 1:500 (WB) |
| Akt1 | Mouse | Invitrogen | AHO1112 | 1:250 (WB) |
| pErk1/2 (pT202/pT204) | Mouse | Invitrogen | 501128621 | 1:1000 (WB) |
| Erk1/2 | Rabbit | Invitrogen | MA515134 | 1:1000 (WB) |
| pGSK3β (pY216) | Rabbit | Invitrogen | 44604G | 1:1000 (WB) |
| GSK3β | Mouse | Invitrogen | MA515597 | 1:1000 (WB) |
| Methyl-PP2a (mL309) | Mouse | Santa Cruz | sc-81603 (2A10) | 1:200 (WB) |
| pPP2a (pY307) | Rabbit | Invitrogen | PIPA536874 | 1:500 (WB) |
| PP2a | Rabbit | Invitrogen | PA517510 | 1:1000 (WB) |
| VDAC1 | Mouse | Millipore Sigma | MABN504 | 1:2000 (WB) |

Anti-tau antibodies were knockout/knockdown validated using either mouse brains, iPSC-derived neurons or both. Anti-pS422 had a strong, non-specific band at ∼50 kDa, slightly beneath a faint p-tau band, in iPSC-derived neurons but not in mouse brains. The non-specific signal occluded reliable quantification in iNeurons. AT100 (pT212/pS214) was not detected in iNeurons, and in concurrence anti-pS214 (Invitrogen, #44742G) was inconsistently detected as a very faint band that could not be reliably quantified.

### Statistical analysis

All experiments were performed using reagents that were aliquoted to minimize freeze-thaw cycles and improve reproducibility. Each ‘n’ was collected independently, as indicated in the figure legends. SC randomized and blinded NNN to sample identifies (lentivirus identities, mouse ages, sample treatments) for data analysis. All analyses were performed using PyCharm Community Edition 2023.2.4. Statistical significance for comparisons between two sample groups was determined by two-tailed Student’s *t* test. Significance for curve comparisons for nocodazole-treated sample groups was determined by post-hoc Tukey’s HSD test. Significance for curve comparisons for MT polymerization reactions was determined by two-way ANOVA analysis.

## ACKNOWLEDGEMENTS

This work was supported by funding to N.N.N. (F31NS098623, F32AG079537, K99AG084855, AARF-22-924146), to E.R. (RF1AG053951), to O.S. (DP2-GM137416, Target ALS, Chan-Zuckerberg Institute), to M.S. (R01AG083949, R01NS095988, R01AG052505), to N.L.K. (P30CA060553 and P41GM108569), to E.B.L. (P30AG072979, P01AG066597) and to Virginia M. Lee (R01AG076434).

## AUTHOR INFORMATION

### AUTHORS AND AFFILIATIONS

**Department of Chemistry, University of Pennsylvania, Philadelphia, PA, USA**

Nima N Naseri, Ibrahim Saleh, Elizabeth Rhoades

**Center for Cellular and Molecular Therapeutics, Children’s Hospital of Philadelphia, Philadelphia, PA, USA**

Nima N Naseri, Sapanna Chantarawong, Ophir Shalem

**Department of Chemistry, Northwestern University, Evanston, Illinois 60208, United States**

Tian Xu, Steven M Patrie, Neil L Kelleher

**Appel Institute for Alzheimer’s Research, and Feil Family Brain & Mind Research Institute, Weill Cornell Medicine, New York, NY, USA**

Manu Sharma

**Department of Pathology and Laboratory Medicine, Perelman School of Medicine, University of Pennsylvania, Philadelphia, PA, USA**

Edward B Lee, Hong Xu

**Department of Genetics, Perelman School of Medicine, University of Pennsylvania, Philadelphia, PA, USA**

Ophir Shalem

**Department of Biochemistry & Biophysics, Perelman School of Medicine, University of Pennsylvania, Philadelphia, PA, USA**

Elizabeth Rhoades

### CONTRIBUTIONS

NNN and ER conceived the project, directed by ER and OS. All experiments were performed and analyzed by NNN unless otherwise indicated. TX performed top-down mass spectrometry experiments under guidance by SMP and NLK. IS generated recombinant 3R tau seeds and performed TEM imaging. We thank Polina Holubovska for providing recombinant 0N3R tau for generating recombinant seeds. SC provided technical support. MS provided mice. EBL and HX provided AD-derived tau seeds and seeding protocols.

## ETHICS DECLARATIONS

The authors declare no competing interests.

## DATA AVAILABILITY

The raw immunoblots generated in this study are available in the supporting information.

