## Supplemental Figures for "Tau aggregation results in an impaired ability to counteract microtubule destabilization via specific combinatorial tau phosphorylation patterns"

### SUPPLEMENTAL FIGURES + FIGURE LEGENDS

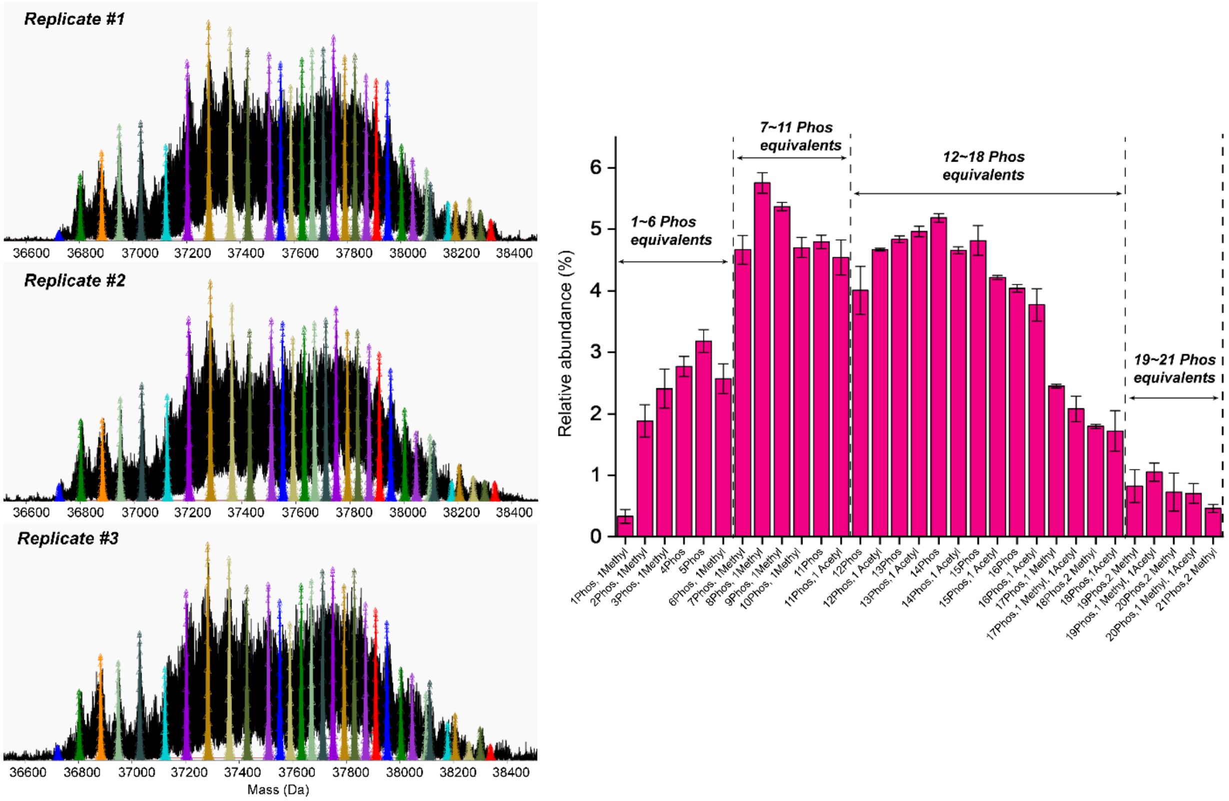


**Figure S1 – Replicate measurements for top-down mass spectrometry analysis of endogenous tau**

Pan-tau was immunoprecipitated from iNeurons (*DPI 30*) as described in Fig. 2a. Three separate replicates for independent I$^2$MS collections are depicted, and annotations are consistently color-coded as in Fig. 2a. The quantification of relative 0N3R tau proteoforms abundance is annotated based on detected PTMs, including phosphorylation (+80 Da), methylation (+14 Da), and acetylation (+42 Da). Quantifications are representative of mean ± STD for n=3 independent I$^2$MS analyses.

#### SUPPLEMENTAL FIGURES + FIGURE LEGENDS

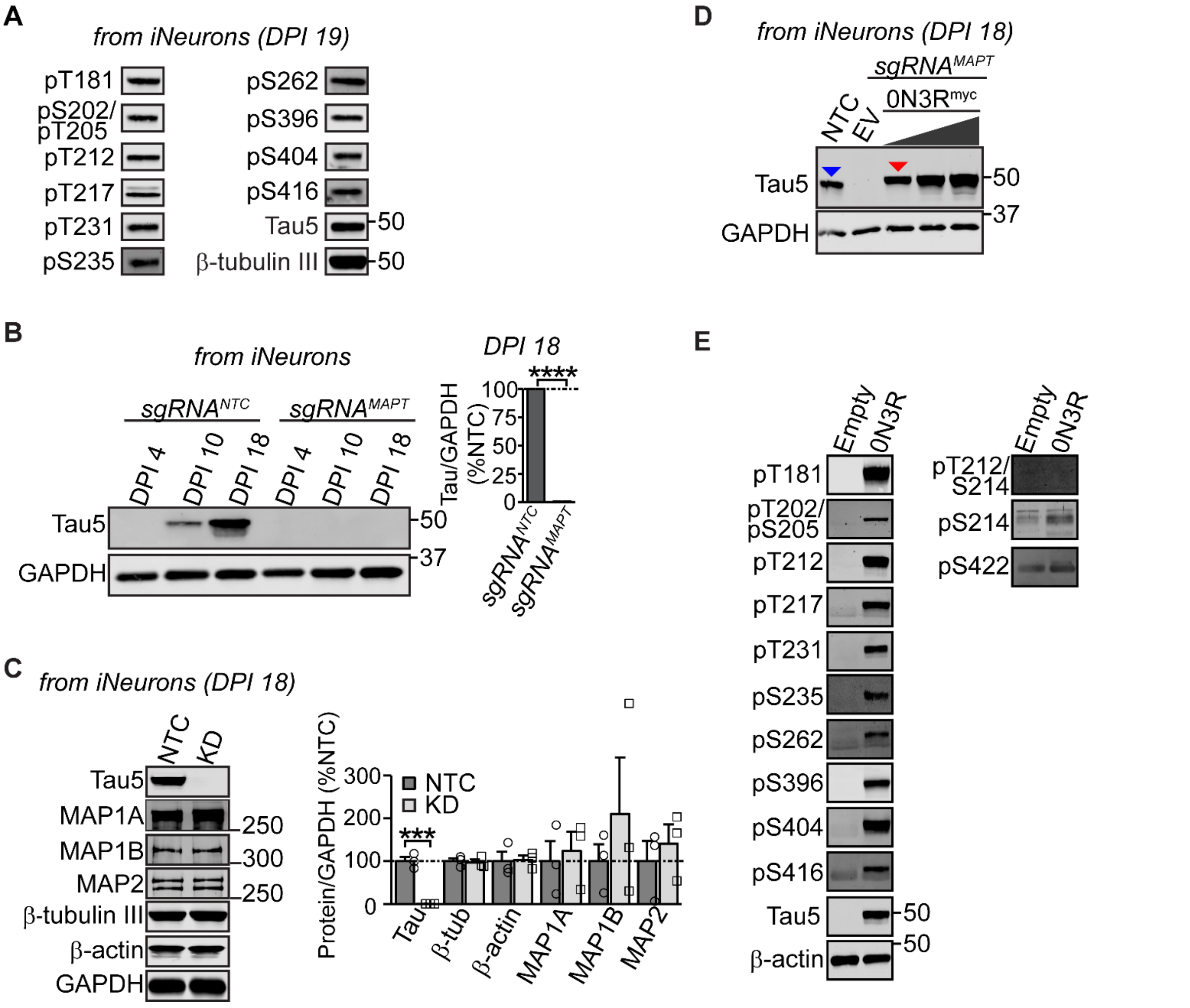


**Figure S2 – iNeuron model generation.**

**a)** P-tau was detected by immunoblot at the indicated epitopes in wildtype human iNeurons. Images are representative of numerous blots. For **b-c)** Ngn2_dCas9-KRAB iPSCs (WTC11) were transduced with either an sgRNA against a non-targeting guide (NTC) for control or *MAPT* to generate tau$^{KD}$ iPSCs. After differentiation, tau levels were undetectable in tau$^{KD}$ iNeurons at 18 days post neuronal-induction (*DPI*), while MAP1A, MAP1B, MAP2, β-tubulin III and β-actin levels were unaffected. N=3 independent cultures and lentiviral transductions per group. **d)** Tau$^{KD}$ neuronal progenitors were lentivirally transduced at *DPI 3*, upon sub-plating, with increasing doses of either empty virus (EV) or 0N3R$^{myc}$ and harvested 9 days later at *DPI 12* for pan-tau (anti-Tau5) detection by immunoblot. The red arrowheads indicate the selected viral titer that was approximately equivalent to endogenous tau expression levels (blue arrowhead). **e)** Tau$^{KD}$ neuronal progenitors were lentivirally transduced as in 'Fig. S2d' with either empty virus (control) or 0N3R$^{myc}$ and harvested at *DPI 12* for p-tau detection by immunoblot. Images are representative of n=3 independent cultures and lentiviral transductions. Phosphorylation was not detected at sites S214, T212/S214 and S422 in iNeurons. ***P<0.001; ****P<0.0001 by Student's t-test. All data are representative of mean ± SEM.

### SUPPLEMENTAL FIGURES + FIGURE LEGENDS

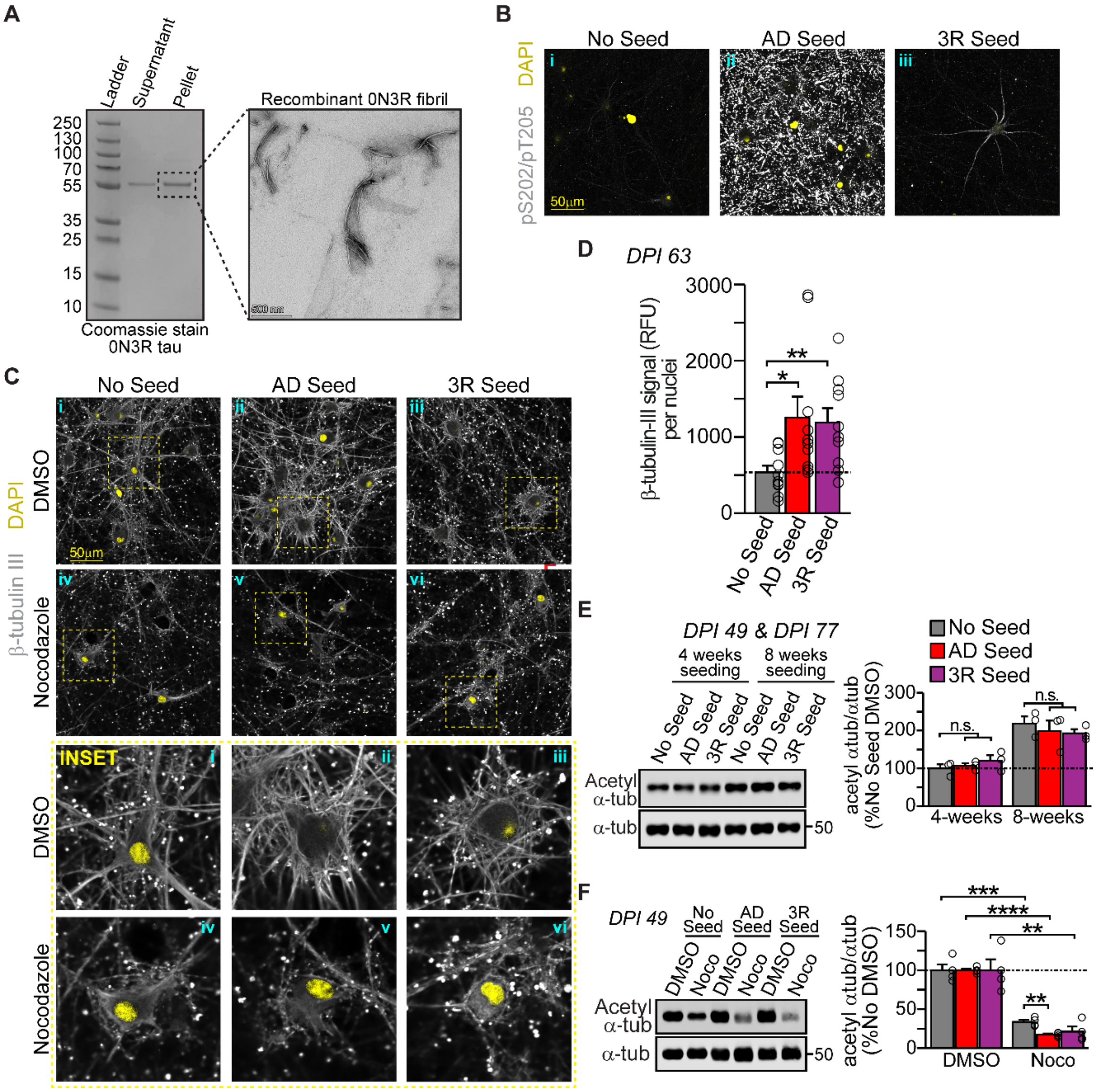

#### SUPPLEMENTAL FIGURES + FIGURE LEGENDS

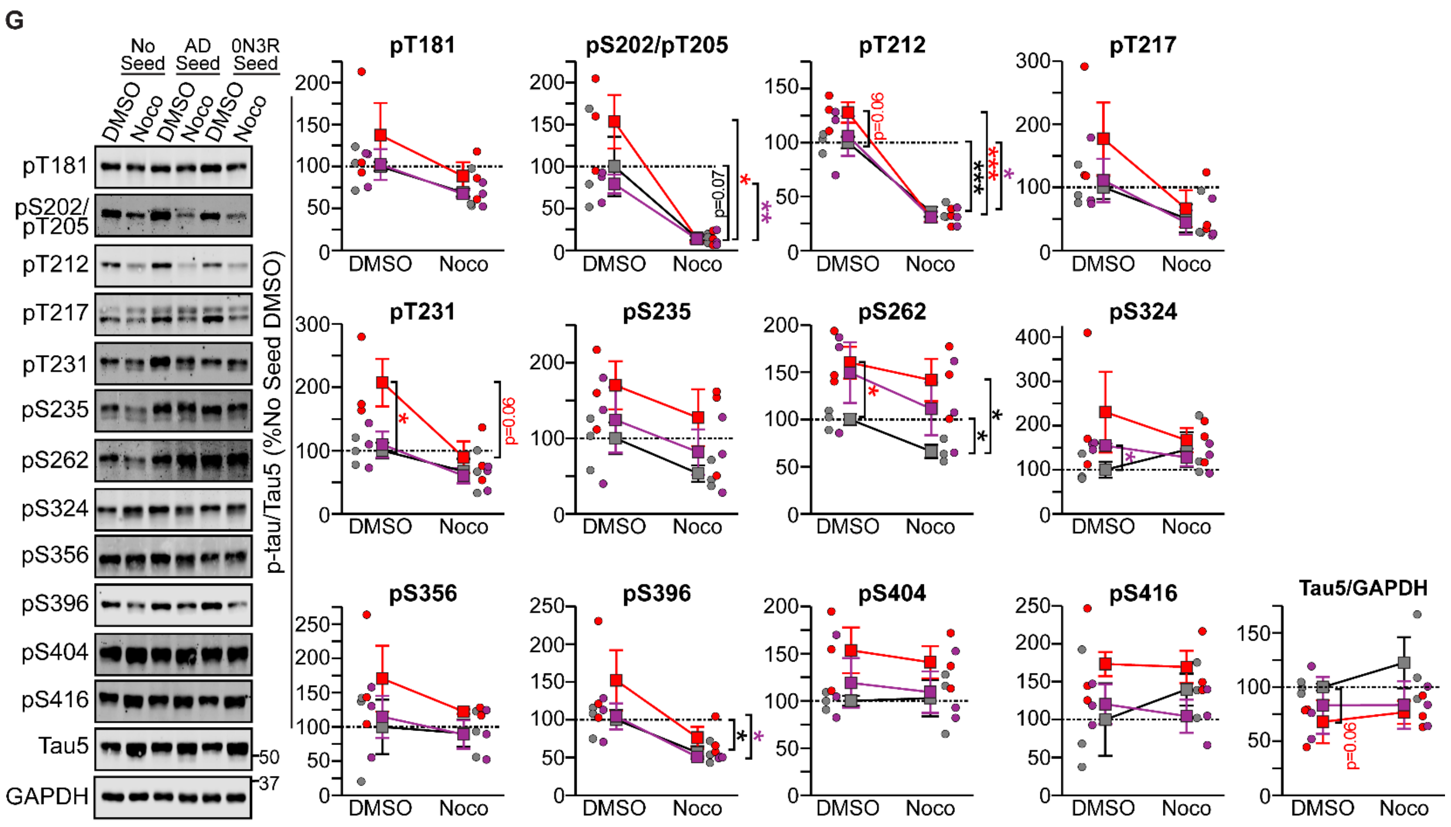


**Figure S3 – Tau seeds induce robust pathology in iNeurons.**

**a)** 50µM recombinant 0N3R was agitated at 1000 RPM for 6 days, followed by centrifugation at 17,000*g* for 90 minutes. The fibrils in the insoluble pellet were imaged by transmission electron microscopy. For **b-c)** As in 'Figure 2c', iNeurons were treated at 3 weeks post-differentiation with insoluble tau seeds either derived from an AD brain or generated from recombinant insoluble 0N3R tau. Soluble proteins were extracted from iNeurons by ice-cold methanol fixation at 6 weeks post-seeding. **b)** Immunocytochemistry was performed against anti-AT8 (pS202/pT205) at 60x magnification. **c)** Immunocytochemistry was performed against anti-β-tubulin III. Images for 'no seed' (panel i) and 'AD seed' (panel ii) were reproduced from 'Fig. 2d'. **d)** Quantification of β-tubulin III signal intensity from 'Fig. 2c' was performed using the mean signal intensity normalized to the number of nuclei in each field of view, from 10 random fields of view, for each sample group. Quantifications for 'no seed' and 'AD seed' were reproduced from 'Fig. 2d'. **e)** Immunoblot and quantification of K40-acetylated α-tubulin normalized to α-tubulin 1A levels at 4-weeks (*DPI 49*) or 8-weeks (*DPI 77*) post-seeding. Blots and quantifications are representative of n=3 independent seeding and treatment experiments. Blots and quantifications for 'no seed' and 'AD seed' were reproduced from 'Fig. 2e'. For **f-g**, unseeded and seeded iNeurons were treated at 4 weeks post-seeding with either DMSO (control) or 10 µM nocodazole for 90 minutes, then harvested in RIPA buffer for Western blot. Blots and quantifications for 'no seed' and 'AD seed' were reproduced from 'Fig. 2f,g'). **f)** Immunoblot and quantification of K40-acetylated α-tubulin normalized to α-tubulin 1A levels. Blots and quantifications are representative of n=4 independent seeding and treatment experiments. **g)** Immunoblot against tau phosphorylated at the indicated phospho-epitopes. Blots and quantifications represent p-tau signal normalized to pan-tau for n=3 independent seeding and treatment experiments. Gray = no seed. Red = AD seed. Purple = 3R seed. All data are representative of mean ± SEM. *P<0.05; **P<0.01; ***P<0.001; ****P<0.0001 by Student's t-test. N.S. = not significant.

#### SUPPLEMENTAL FIGURES + FIGURE LEGENDS

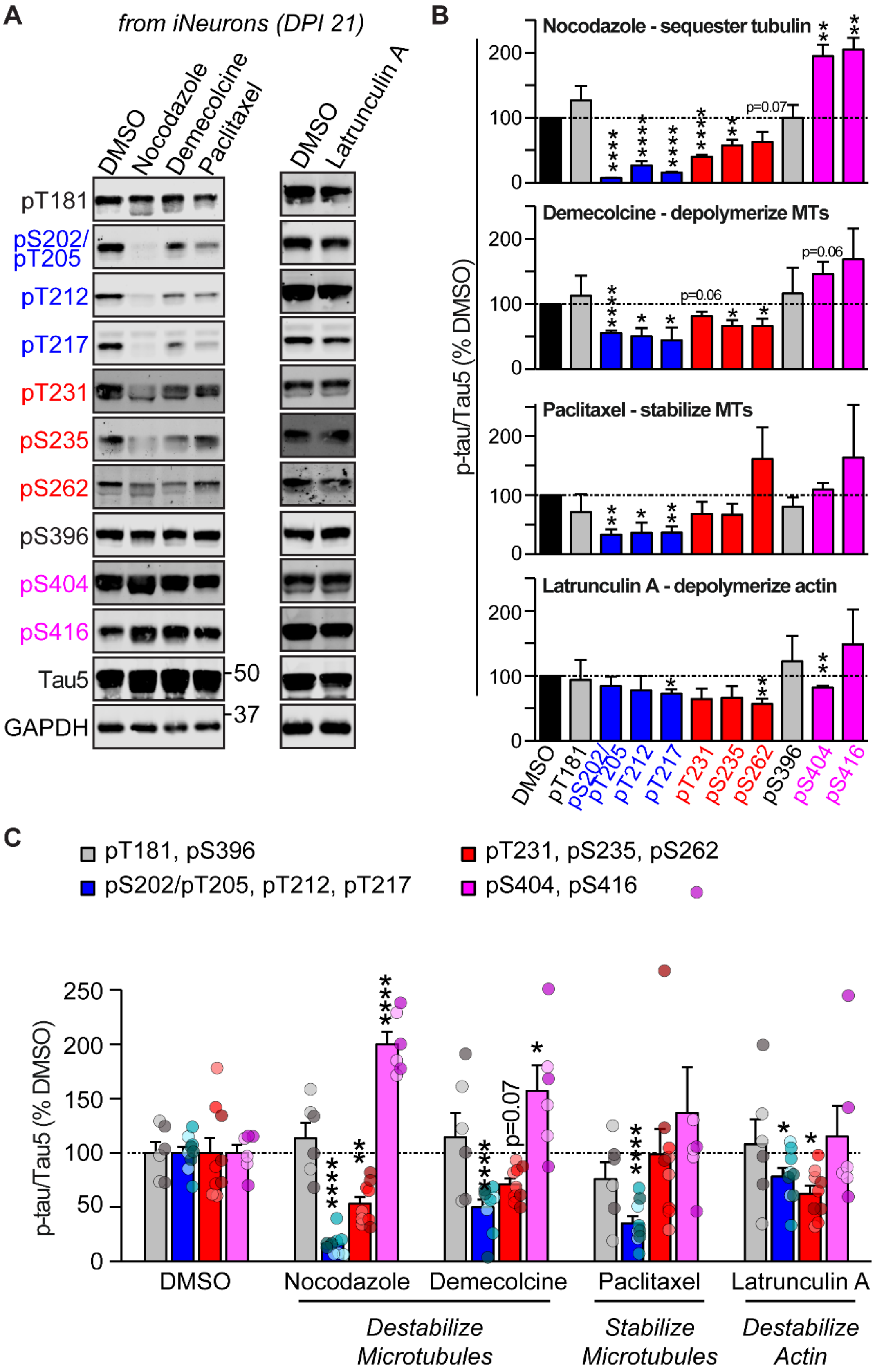


**Figure S4 – Tau phosphorylation is primarily influenced by alterations to MT stability.**

**a)** iNeurons (*DPI 21*) were treated with either DMSO, nocodazole (10 µM), demecolcine (1 µM), paclitaxel (10 µM) or

### SUPPLEMENTAL FIGURES + FIGURE LEGENDS

latrunculin A (2 μM) for 90 minutes to manipulate the cytoskeleton. Samples were separated by SDS-PAGE followed by anti-p-tau immunoblot. P-tau levels were quantified and normalized to pan-tau (anti-Tau5) for each treatment and then normalized to DMSO-treated samples (control). N=3 independent cultures and treatments per group. **b)** Quantification of 'Fig. S4a'. P-tau signals were normalized to pan-tau (Tau5) and plotted as a percentage of DMSO levels for each manipulation. **c)** Consolidated, grouped quantifications of 'Fig. S4a'. Tau phosphorylation was primarily influenced by alterations to MT stability. Actin destabilization may indirectly impact MT stability, and thus may exert some indirect effects on tau phosphorylation at certain epitopes. All data are representative of mean ± SEM. *P<0.05; **P<0.01; ****P<0.0001 by Student's t-test.

#### SUPPLEMENTAL FIGURES + FIGURE LEGENDS

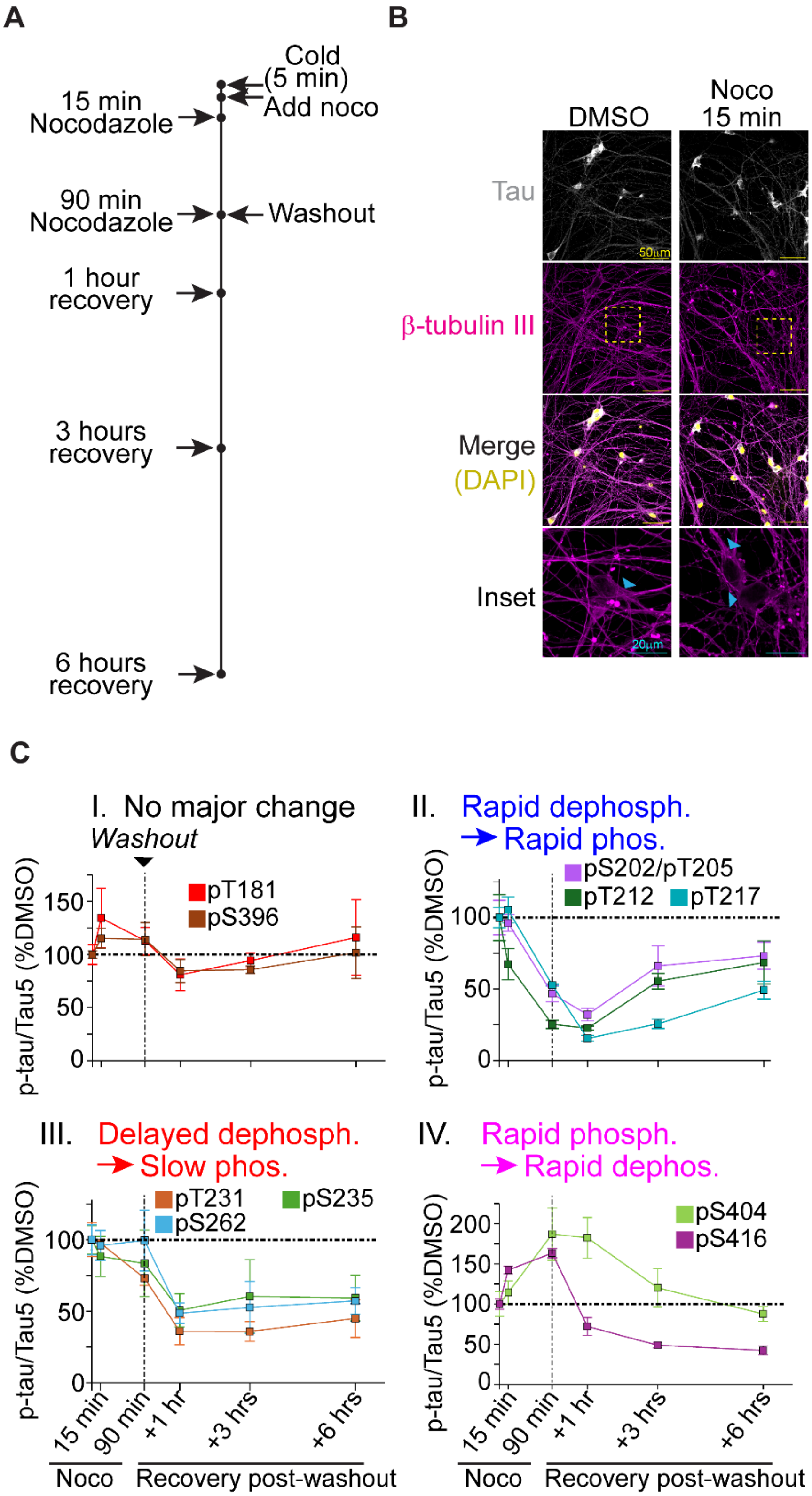


**Figure S5 – Individual phospho-epitope results for nocodazole time course.**

**a)** Treatment paradigm for nocodazole treatment and washout in iNeurons. **b)** Nocodazole treatment disrupted tau-MT networks in iNeurons, as visualized by anti-tau and anti-β-tubulin III immunofluorescence. **c)** Site-specific changes to tau phosphorylation during the time course assay, as described in 'Fig. 3c'.

#### SUPPLEMENTAL FIGURES + FIGURE LEGENDS

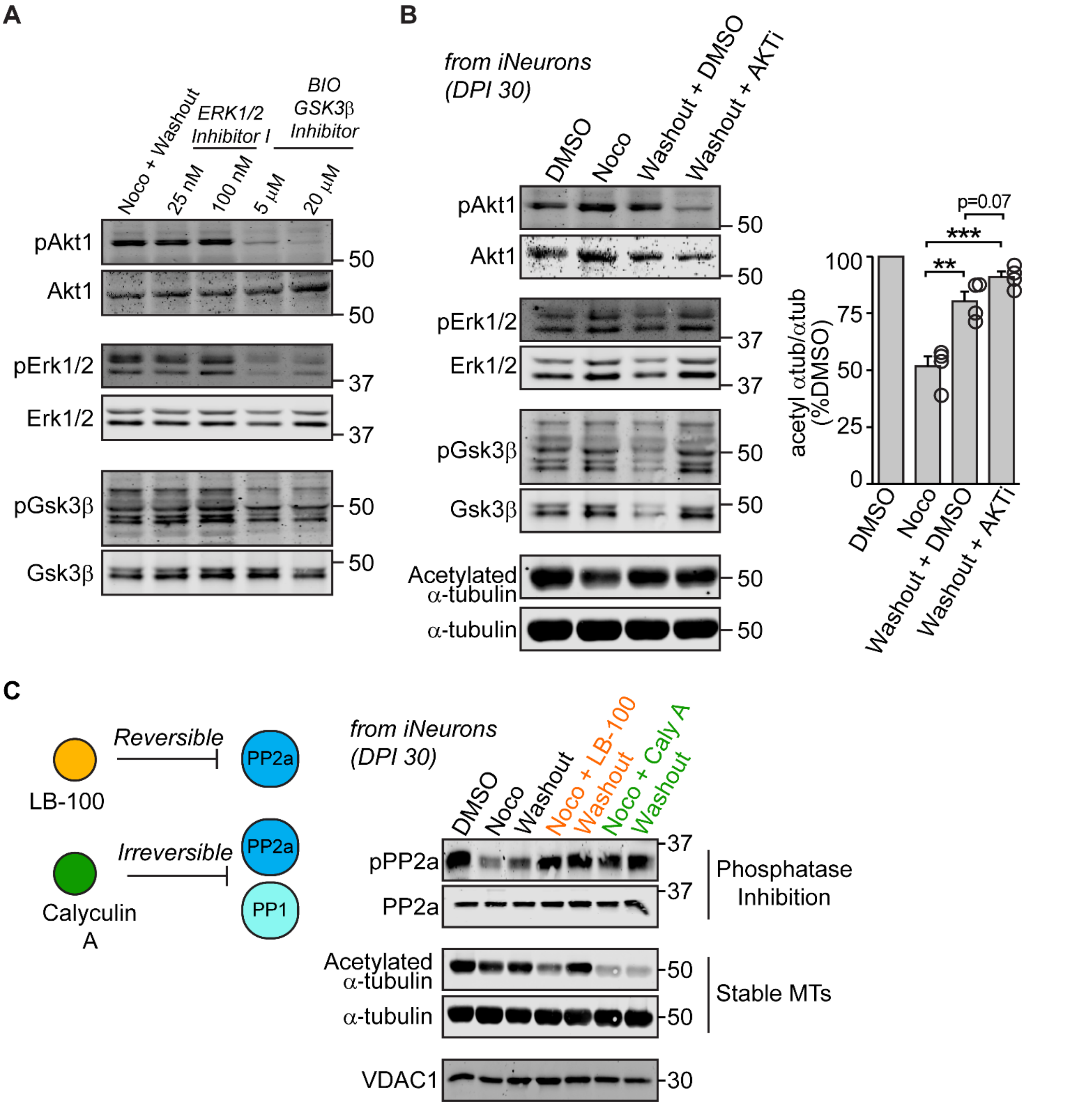


**Figure S6 – PP2a phosphatase inhibition but not Akt1 kinase inhibition significantly impacted MT recovery in response to nocodazole treatment.**

**a)** iNeurons (*DPI 30*) were treated with 10 μM nocodazole for 90 minutes followed by washout for 3 hours. After nocodazole washout, iNeurons were treated for 3 hours with either DMSO, ERK1/2 Inhibitor I (25 nM or 100 nM) or BIO GSK3β Inhibitor (5 μM or 20 μM). The neurons were harvested for SDS-PAGE followed by immunoblot against markers for kinase activation: phosphorylation against Akt1 (S473), Erk1/2 (T202/T204) and GSK3β (Y216). Blots are representative of n=3 independent experiments. **b)** iNeurons (*DPI 30*) were treated with nocodazole (10 μM) for 90 minutes, followed by nocodazole washout for three hours. AKT1 inhibitor VIII (20 μM) was supplemented during the washout period. Neurons were then harvested for immunoblot to determine successful and specific Akt1 kinase inhibition, as well as changes to MT stability as measured by acetylated-α-tubulin (K40) normalized to total α-tubulin levels (as a percentage of DMSO treated controls). N=4 independent treatments. For **c-d)** iNeurons (*DPI 30*) were simultaneously treated with nocodazole and the indicated PP2a inhibitors (LB-100 at 20 μM; Calyculin A at 20 nM). The nocodazole and phosphatase inhibitors were washed out and neurons were allowed to recover for 3 hours. Successful PP2a inhibition was determined by anti-phosho-PP2a (Y307) immunoblot. Changes to MT stability were determined by anti-acetylated-α-tubulin (K40) immunoblot. Blots are representative of n=4 independent treatments. **P<0.01; ***P<0.001 by Student's t-test. All data are representative of mean ± SEM.

#### SUPPLEMENTAL FIGURES + FIGURE LEGENDS

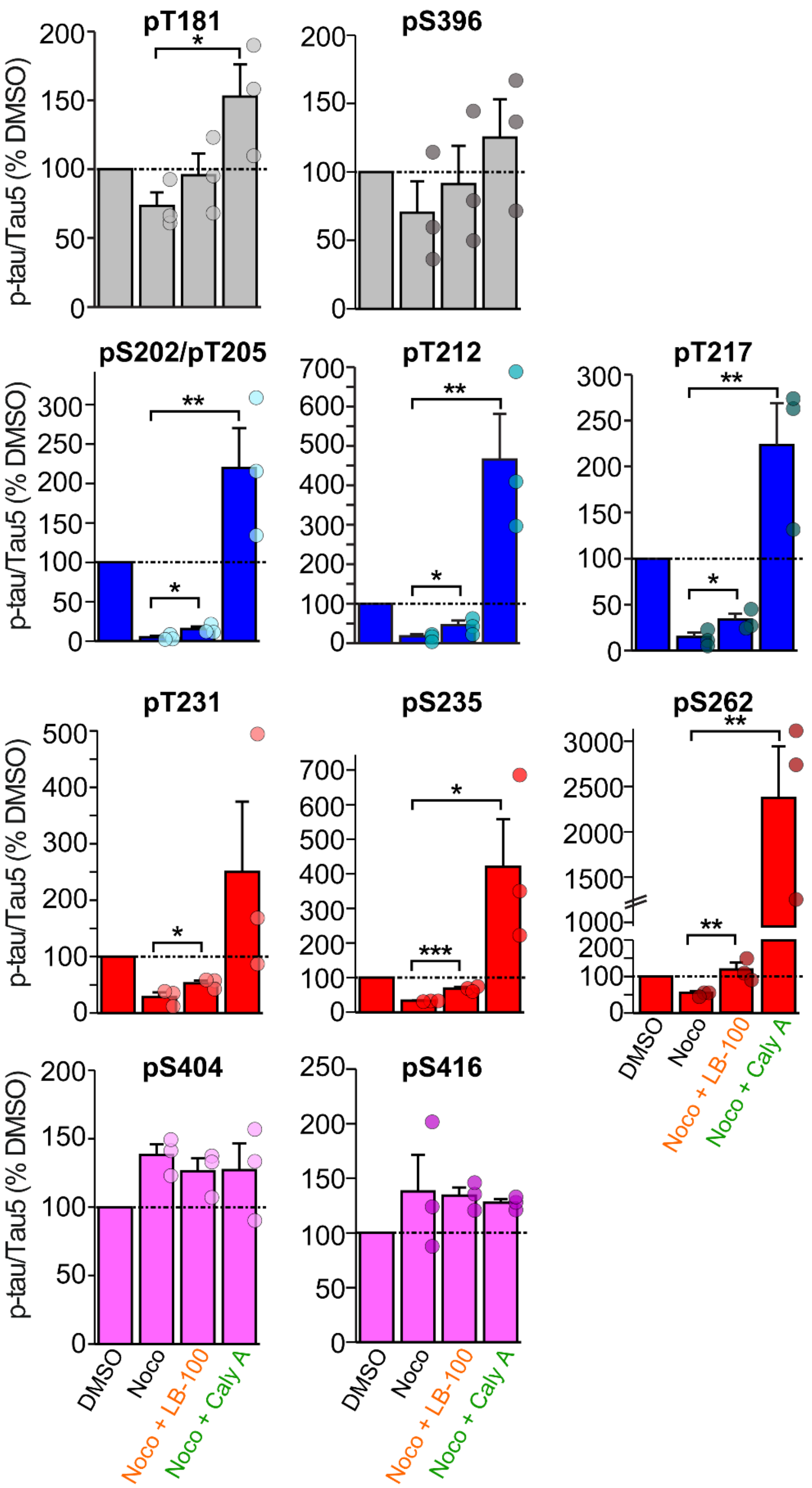


**Figure S7 – Individual phospho-epitope results for phosphatase inhibition during nocodazole-induced MT catastrophe.**

As described in 'Fig. 4c-e', iNeurons were simultaneously treated with nocodazole (10 μM) and either LB-100 (20 μM) or Calyculin A (20 nM) for 90 minutes. P-tau levels were normalized to total tau (anti-Tau5) for each sample, then quantified

### SUPPLEMENTAL FIGURES + FIGURE LEGENDS

as a percentage of DMSO-treated (control) samples. N=3 independent cultures and treatments per group. *P<0.05; **P<0.01 by Student’s t-test. All data are representative of mean ± SEM.

#### SUPPLEMENTAL FIGURES + FIGURE LEGENDS

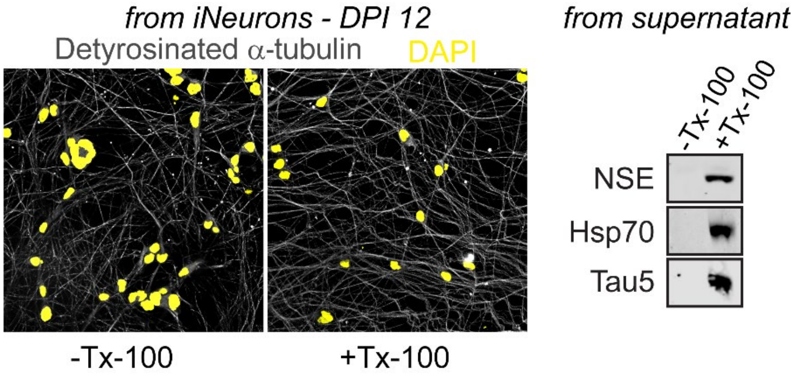


**Figure S8 –MT fractionation in iNeurons.**

Soluble tubulin was extracted by 1% Triton-X 100 in MT stabilization buffer and the remaining, intact MTs remained adhered to the culture dish. The remaining intact MTs were visualized by anti-detyrosinated α-tubulin, a marker for stable, polymerized MTs. Cytosolic proteins such as NSE, Hsp70 and tau were leaked out of the neurons and collected from the culture supernatant

#### SUPPLEMENTAL FIGURES + FIGURE LEGENDS

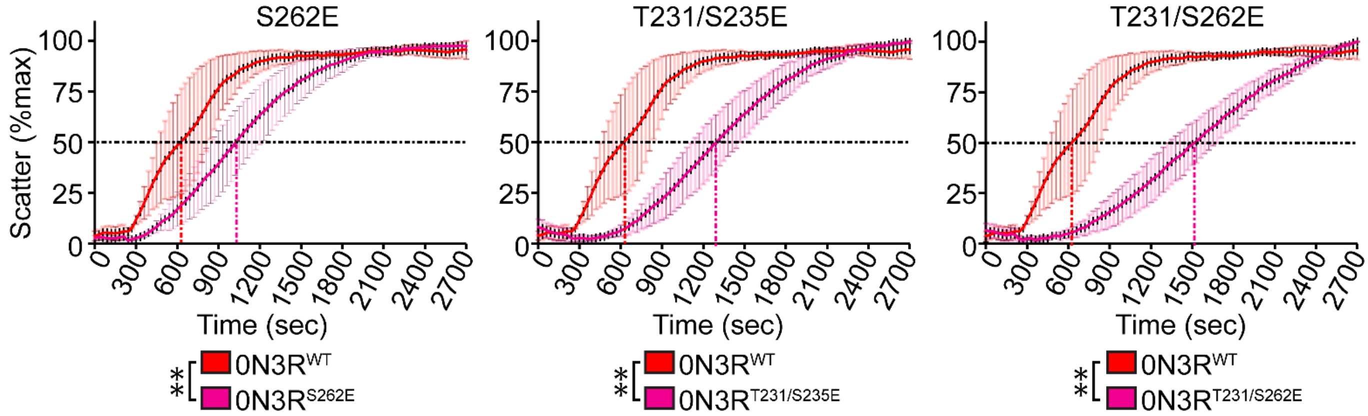


**Figure S9 – Isoform-dependent effects of phospho-mimics on tau's ability to promote MT polymerization.**

MT polymerization curves by light scatter assay for phospho-mimics S262E, T231E/S235E and T231E/S262E, which all exhibited strong effects on tau's ability to promote MT polymerization, as described in Fig. 5c. All data are representative of mean ± SEM for 3 independent polymerization reactions. N.S. = not significant. *P<0.05; **P<0.01 by two-way ANOVA.

### SUPPLEMENTAL FIGURES + FIGURE LEGENDS

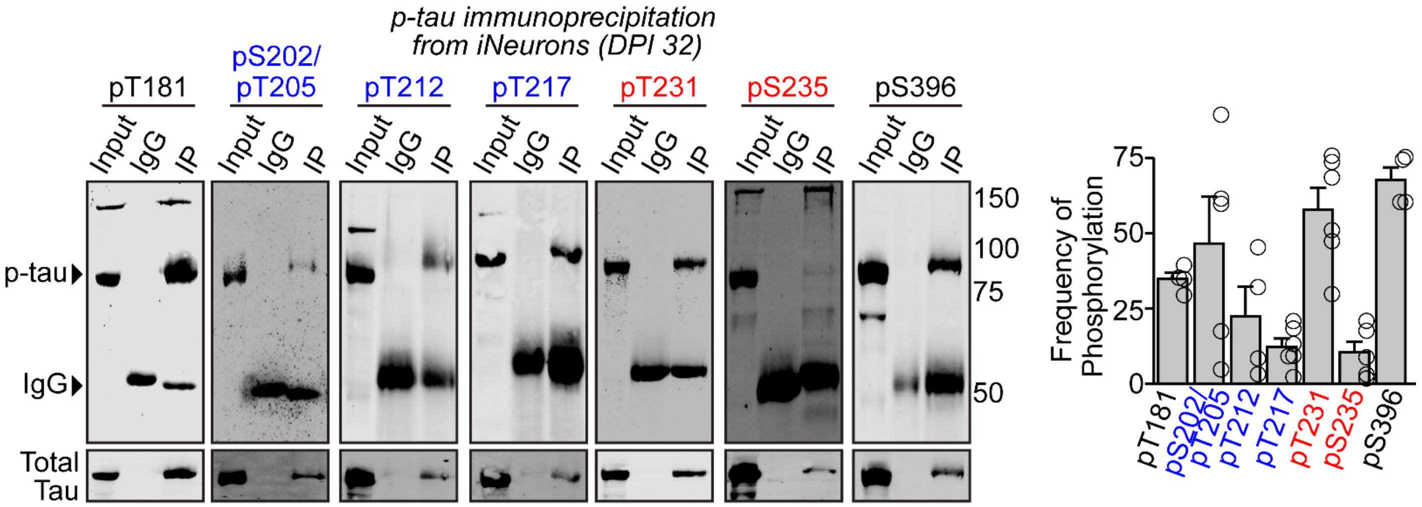


$$\text{Phospho-Occupancy} = \frac{\text{signal for efficiency of captured pan-tau}}{\text{affinity capture efficiency of p-tau}} \times 100 = \frac{\frac{\text{pan-tau IP signal}}{\text{pan-tau input signal}}}{\frac{\text{p-tau IP signal}}{\text{p-tau input signal}}} \times 100$$

***e.g., calculations for a single replicate for pS396 immunoprecipitation***

$$\frac{\frac{\text{pan-tau IP signal from pS396 IP}}{\text{pan-tau input from pS396 IP}}}{\frac{\text{p-tau IP signal from pS396 IP}}{\text{p-tau input signal from pS396 IP}}} \times 100 = \frac{\frac{1.12\text{x}10^5}{2.33\text{x}10^5 * \cancel{44.7}}}{\frac{4.28\text{x}10^5}{5.28\text{x}10^5 * \cancel{44.7}}} \times 100 = \frac{0.481}{0.811} \times 100 = \text{59.3\% of captured total tau was positive for pS396}$$

*dil. factor*

**Figure S10: Tau phosphorylation occupancy levels at various epitopes in iNeurons.**

$Tau^{KD}$ iNeurons were lentivirally transduced with $0N3R^{EGFP}$, and p-tau was then immunoprecipitated using the indicated p-tau antibody, separated by SDS-PAGE and immunoblotted against the indicated immunoprecipitated p-tau epitope and pan-tau (anti-Tau5 mouse for p-tau rabbit immunoprecipitations; anti-Tau5 rabbit for p-tau mouse immunoprecipitations). IgG control = anti-myc mouse monoclonal or anti-GAPDH rabbit polyclonal antibodies. Quantification of phospho-site abundance, described in the 'Methods' section, is representative of the average of two technical replicates for n=4 (pT181, pT212, pS396), n=5 (AT8) and n=6 (pT217, pT231, pS235) independent cultures and immunoprecipitations. An example of the back-calculation method is provided for a single replicate (out of 4 replicates) for quantification of phosphate occupancy at S396. All data are representative of mean ± SEM.
